# TIM-3 blockade enhances ex vivo stimulated allogeneic NK cell therapy for relapsed murine neuroblastoma after hematopoietic cell transplant

**DOI:** 10.1101/2024.07.09.602731

**Authors:** Aicha E Quamine, Nicholas R Mohrdieck, Chloe A King, Anastasia A Griggs, Jillian M Kline, Monica M Cho, Sean P Rinella, Katharine E Tippins, Paul D Bates, Lei Shi, Longzhen Song, Nicholas J Hess, Tyce J Kearl, Bryon D Johnson, Christian M Capitini

## Abstract

**Background:** High-risk neuroblastoma (HR-NBL) is an aggressive tumor of the sympathetic nervous system with high risk of relapse and poor overall survival. Allogeneic hematopoietic cell transplant (allo-HCT) has been used previously in HR-NBL patients; however, graft-versus-host-disease (GVHD) and disease progression have limited clinical application. Ex-vivo stimulated allogeneic natural killer (NK) cells represent a potential approach to enhance the graft-versus-tumor (GVT) effect without exacerbation of GVHD but have not shown efficacy in NBL.

**Methods:** Ex-vivo stimulated NK cells from C57BL/6NCr (B6) mice were expanded with soluble IL-15/IL-15Rα alone or with irradiated CD137L/CD54^+^ AgN2a-4P (15-4P) at a 1:1 ratio for 10-12 days. Allogeneic NK cells were then analyzed for activation, proliferation, cytokine production, and cytotoxicity against two murine NBL cell lines, Neuro2a and NXS2, in the absence or presence of anti-TIM-3. Lethally irradiated B6AJF1 Mice received allo-HCT from B6 donors followed by NBL challenge after 7 days to mimic tumor relapse. Select groups received anti-TIM-3 starting on day 9 for every 4 days with/without infusions of 15-4P B6 NK cells on days 14, 21, and 28. In select experiments, T cell and NK cells were selectively depleted to establish their contribution to the GVT effect. All groups were analyzed for tumor growth, GVHD and overall survival.

**Results:** Co-culturing NK cells with 15-4P results in 78-fold expansion with increased expression of Ki-67 and NKG2D, NKp46, TRAIL and TIM-3. 15-4P stimulated allogeneic NK cells showed enhanced cytotoxicity against NBL compared to IL-15 NK cells alone but was limited in part due to high expression of TIM-3 ligands on Neuro-2a compared to NXS2. The addition of TIM-3 blockade further enhanced NK cytotoxicity versus Neuro-2a, with enhanced 15-4P NK cell degranulation, Eomes, TRAIL and FasL expression observed. Analysis of RNA from 15-4P NK cells exposed to TIM-3 blockade showed gene expression of chemokines, NKG2D/DAP12 signaling, non-canonical NF-κb pathway and TRAIL signaling. Blockade of NKG2D, TRAIL or FasL on 15-4P NK cells abrogated cytotoxicity. In vivo, the combination of 15-4P stimulated allogeneic NK cells and TIM-3 blockade after allo-HCT resulted in prolonged survival against NBL with decreased tumor burden compared to NK cells or anti-TIM-3 alone, without inducing GVHD. Depletion of NK cells, but not T cells, abrogated the GVT effect.

**Conclusion:** Allo-HCT can be a platform for treating NBL using combination ex-vivo stimulated allogeneic NK cell therapy with TIM-3 blockade to enhance the GVT effect without inducing GVHD.

**Ethics Statement:** The animal study M005915 was reviewed and approved by University of Wisconsin-Madison IACUC.

**WHAT IS ALREADY KNOWN ON THIS TOPIC:**

- T cell depleted (TCD) allogeneic hematopoietic cells transplant (allo-HCT) has potential to be a salvage therapy for relapsed/refractory neuroblastoma (NBL) through the graft-versus-tumor (GVT) effect, however, the high incidence of graft-vs-host disease (GVHD) and disease progression has hindered widespread clinical application. Allogeneic NK cell therapy can be a safe and feasible treatment but has had limited efficacy in NBL. TIM-3 blockade has shown encouraging results for a variety of tumors but has not been explored for NBL nor used to enhance the GVT effect.

**WHAT THIS STUDY ADDS:**

- Ex vivo stimulated allogeneic NK cells demonstrate robust expansion, proliferation, activation and cytotoxicity against NBL which is further augmented by the addition of TIM-3 checkpoint blockade. In an allo-HCT model of NBL relapse, mice treated with IL-15/IL-15Rα/AgN2a-4P (15-4P) stimulated NK cells and anti-TIM-3 exhibited prolonged overall survival and reduced tumor growth without exacerbating GVHD.

**HOW THIS STUDY MIGHT AFFECT RESEARCH, PRACTICE OR POLICY:**

- Relapse of NBL after TCD allo-HCT can be successfully treated with the combination of adoptively transferred 15-4P stimulated allogeneic NK cells and TIM-3 checkpoint blockade, representing a novel therapeutic approach that can be translated to the clinic or used as a platform to treat other solid tumors with high TIM-3 ligand expression.

## INTRODUCTION

High-risk neuroblastoma (NBL) is an aggressive extracranial solid tumor of neural crest origin that commonly occurs in children typically under 5 years old. About 40% of NBL patients are considered high risk due to advanced age, unfavorable tumor characteristics such as low burden of recurrent somatic mutations, MYCN amplification, and metastasis to sites such as the bone, liver and regional lymph nodes [1]. These molecular characteristics indicate aggressive disease and poor prognosis which necessitates intensive treatment regimens including chemotherapy, surgery, consolidation with autologous hematopoietic cell transplant (auto-HCT), radiation therapy and post-consolidation anti-GD2 immunotherapy with GM-CSF and cis-retinoic acid. The addition of anti-GD2 therapy has increased the 5-year overall survival of high-risk NBL from 20 to 50 percent, yet half of patients still relapse [2]. There is still a critical need for the development of therapies for NBL with lower toxicities [3].

Auto-HCT has been standard of care for high-risk NBL for decades but offers little benefit outside of stem cell rescue. Three phase III randomized clinical trials have reported that supratherapeutic chemotherapy with auto-HCT support improved event free survival (EFS) but failed to ameliorate 5-year overall survival (OS) which highlights a critical need for incorporating novel therapies for the treatment of high-risk NBL [4,5]. Allogeneic HCT (allo-HCT) offers a unique advantage to auto-HCT by leveraging donor lymphocyte-mediated eradiation of residual disease. However, it has been difficult to translate these results to solid tumors due to the high incidence of graft-vs-host disease (GVHD) and disease progression [6].

Advances in graft engineering now allow for precise removal of allogeneic T cells (e.g. αβ-TCR depletion, CD3 depletion, etc.) that can greatly reduce the risk of GVHD, however these methods increase likelihood of relapse, infection, and delays immune reconstitution [7,8]. Adoptive cell transfer of natural killer (NK) cells after T cell depleted allo-HCT has the potential to improve outcomes by treating NBL relapse while controlling viral infections without inducing GVHD. NK cells are innate lymphoid cells that play an important role in anti-tumor surveillance by identifying infected, stressed, or tumorigenic cells through a balance of activating and inhibitory cell surface receptors [9,10]. NK cells are not limited by antigen recognition, instead missing or mismatched binding between killer-cell-immunoglobulin (KIR) and “non-self” major histocompatibility complex (MHC) class I (or human leukocyte antigens (HLA) class I) triggers a cytotoxic response [11]. To evade T cell recognition, NBL cells downregulate MHC class I which exposes them to NK cell mediated killing. NK cells do not induce GVHD and can exert a protective defense against GVHD by suppressing allo-reactive T cells, making them a safe option for an off-the-shelf therapy [12]. Clinical studies in NBL have confirmed the safety profile of allogeneic NK cell therapy, however response rates remain modest. Due to the biological limitations of naïve NK cells, methods of ex vivo expansion and activation that enhance NK cell cytotoxicity without inducing exhaustion are critical.

Expression of TIM-3 on NK cells has been linked to both inhibitory and activating roles in a pathway-dependent manner. Although increased TIM-3 expression correlates with T cell dysfunction, NK cells with high TIM-3 expression are highly responsive and robustly degranulate against target cells and TIM-3 expression increases following cytokine stimulation and promotes IFN-γ production [13]. Cytokine stimulation with multiple cytokines, including IL-15, have been shown to induce TIM-3 expression on NK cells suggesting that TIM-3 expression marks a mature and highly activated subpopulation [14]. However, cross-linking of TIM-3 with agonist antibodies or cognate ligands significantly inhibits NK cell–mediated cytotoxicity induced by CD16 and NK cell activation and dampens degranulation [15].

Here, we explore the use of AgN2a-4P, a murine neuroblastoma cell line engineered from an aggressive variant of the Neuro-2a murine neuroblastoma cell line (AgN2a) to express four immune costimulatory proteins CD54, CD80, CD86, and CD137L (4P) on its surface. Notably, CD137L (41BBL) serves as a well-established NK cell activator, driving proliferation and heightened antibody-dependent cellular cytotoxicity (ADCC) via Fc receptor signaling [16]. Additionally, CD54 can bind to LFA-1 and MAC-1 on NK cells and augments cytotoxicity by promoting adhesion and granule polarization. The current investigation explores the impact of adoptive cell therapy of ex-vivo AgN2a-4P-stimulated allogeneic NK cells with TIM-3 immune checkpoint blockade for the first time after allo-HCT for NBL as a novel combination immunotherapy.

## METHODS

### Mice

Female C57BL/6NCr (B6, H-2b), and B6AJF1 (B6AJ, H-2b x H-2a) mice were purchased from the National Cancer Institute (NCI) Animal Production Program and Charles River Laboratories International (Frederick, MD). Mice were housed in accordance with the Guide for Care and Use for Laboratory Mice and used between 8 and 16 weeks of age at time of experimentation. The Institutional Animal Care and Use Committees (IACUC) at the University of Wisconsin-Madison (M005915) approved all protocols.

### Murine ex vivo NK isolation and activation

Murine bone marrow cells (BMCs) were harvested from tibias and fibulas and processed into a single cell suspension. Red blood cells were lysed with ACK lysis buffer (Lonza, Walkersville, MD). NK cells were isolated using the NK cell isolation kit (Miltenyi Biotec, Auburn, CA) by negative selection on the AutoMACs Pro (Miltenyi Biotec). NK cells were expanded and activated in the presence or absence of irradiated (100Gy) AgN2a-4P at a 1:1 ratio and cultured in RPMI medium (Corning Life Sciences, Durham, NC) containing 10 ng/ml IL-15/IL-15Rα conjugate, comprised of recombinant mouse IL-15 protein (R&D Systems, Minneapolis, MN) and recombinant mouse IL-15Ralpha Fc chimera protein (R&D Systems), 10% fetal bovine serum (GeminiBio, Sacramento, CA) with 50 IU/mL penicillin/streptomycin (Lonza), 0.11 mM 2-beta-mercaptoethanol (Gibco, Carlsbad, CA), 1X MEM nonessential amino acids (Corning), 25 mM HEPES buffer (Corning), and 1 mM sodium pyruvate (Corning). NK cells were expanded for 12 days and were supplemented with additional 10ng/mL IL-15/IL-15Rα and 5 mL of media every 2 days and maintained at 37°C in 5% CO_2_.

### Allogeneic bone marrow transplant

BMCs were harvested from B6 mice and processed into single cell suspensions. T cells were depleted using a CD3 MicroBead Kit (Miltenyi Biotec) by negative selection on the AutoMACs Pro Separator (Miltenyi Biotec) at day +0. Separately, T cells were isolated using a pan T cell Isolation Kit (Miltenyi Biotec) on single cell suspensions obtained from splenocytes isolated from B6 mice. B6AJF1 recipient mice received a single dose of 11 Gy total body irradiation using an X-Rad 320 (Precision Xray Irradiation, Madison, CT). Irradiated recipients received retroorbital injection of 5 x 10^6^ donor-derived BMCs and 1 x 10^6^ T cells in serum-free RPMI (Invitrogen, Carlsbad, CA).

### In vivo tumor challenge and immunotherapy treatment

At day +7, 2×10^6^ Neuro2A NBL cells (H-2a) were prepared as a single cell suspension in serum-free RPMI and injected into B6AJF1 recipient mice subcutaneously on the right hind limb flank to mimic post-HCT relapse. Tumor length and width was measured biweekly using a digital caliper and tumor growth was calculated as length x width (mm^2^). Mice were euthanized when tumors reached 2 cm in any dimension. Tumor measurements from mice that died prior to the experiment endpoint were carried for the remainder of the study for statistical comparison. Mice that received adoptive NK cell therapy received 2×10^6^ B6 donor-derived NK cells expanded in the presence of IL-15/IL-15Rα and with or without irradiated AgN2a-4P. Expanded NK cells were then harvested after 12 days of expansion and administered by intravenous injection on days +14, +21, and +28. Mice receiving immune checkpoint blockade were given intraperitoneal injection of 250μg of anti-TIM-3 (BioXcell Cat. No. BE0115) every 4 days starting on day +9. Control mice were treated on an identical injection schedule with 250μg of IgG2a isotype control antibody (BioXcell Cat. No. BE0089).

### Statistical analysis

Statistics were performed using GraphPad Prism version 9.0 (GraphPad Software, San Diego, CA). Survival curves were plotted using Kaplan-Meier estimates, and the Mantel-Cox log-rank test was used to analyze survival data. The 2-tailed Mann-Whitney U test (non-parametric data sets) or unpaired T test (parametric data sets) with Welch’s corrections were used for the statistical analysis when comparing 2 groups. When comparing 3 or more groups the Kruskal-Wallis (non-parametric data sets) with Dunn’s multiple comparison post-hoc test or One-way ANOVA (parametric data sets) with Tukey post-hoc test. P < 0.05 was considered statistically significant.

Refer to supplemental methods for more information.

## RESULTS

### AgN2a-4P stimulated ex-vivo activated NK cells show increased expression of activation and proliferation markers

AgN2a-4P is a genetically engineered murine NBL cell line that expresses CD54, CD80, CD86, and CD137L co-stimulatory molecules but has not been tested on allogeneic NK cells previously. AgN2a-4P can interact with NK cells through CD137L (which binds to CD137 on NK cells) and CD54 (which binds to LFA-1/MAC-1 on NK cells) [17] (Fig. 1A). To study the impact of AgN2a-4P exposure on NK cell expansion and activation, NK cells from B6 mice were expanded with AgN2a-4P cells with IL-15/IL-15Rα at a 1:1 ratio. After 12 days of coculture, activation marker expression was assessed by flow cytometry (Fig. 1B). B6 NK cells expanded with IL-15/IL-15Rα and AgN2a-4P (15-4P) exhibited a significant increase in the percentage of NKG2D+ cells (29.27 ± 1.894) compared to IL-15 NK cells (20.80 ± 0.7095). While 15-4P did not impact the percentage of NKp46^+^ NK cells (35.08 ± 1.427) in comparison to IL-15 NK cells (33.70 ± 0.3342), the expression of NKp46 by MFI was significantly increased (p = 0.0286) (Fig. 1C). Due to expression of 41BBL on AgN2a-4P we confirmed whether AgN2a-4P exposure during expansion impacts NK cell proliferation. We observed 78-fold expansion in 15-4P (78.33 ± 20.46) compared to IL-15 NK cells (27.39 ± 1.207) (Fig. 1D). There was a significant increase (p = 0.0006) of Ki-67^+^ 15-4P NK cells (52.24 ± 3.953) and Ki-67 expression by MFI compared to IL-15 NK cells (13.42 ± 0.318) (Fig. 1E-F).

**Figure 1.**
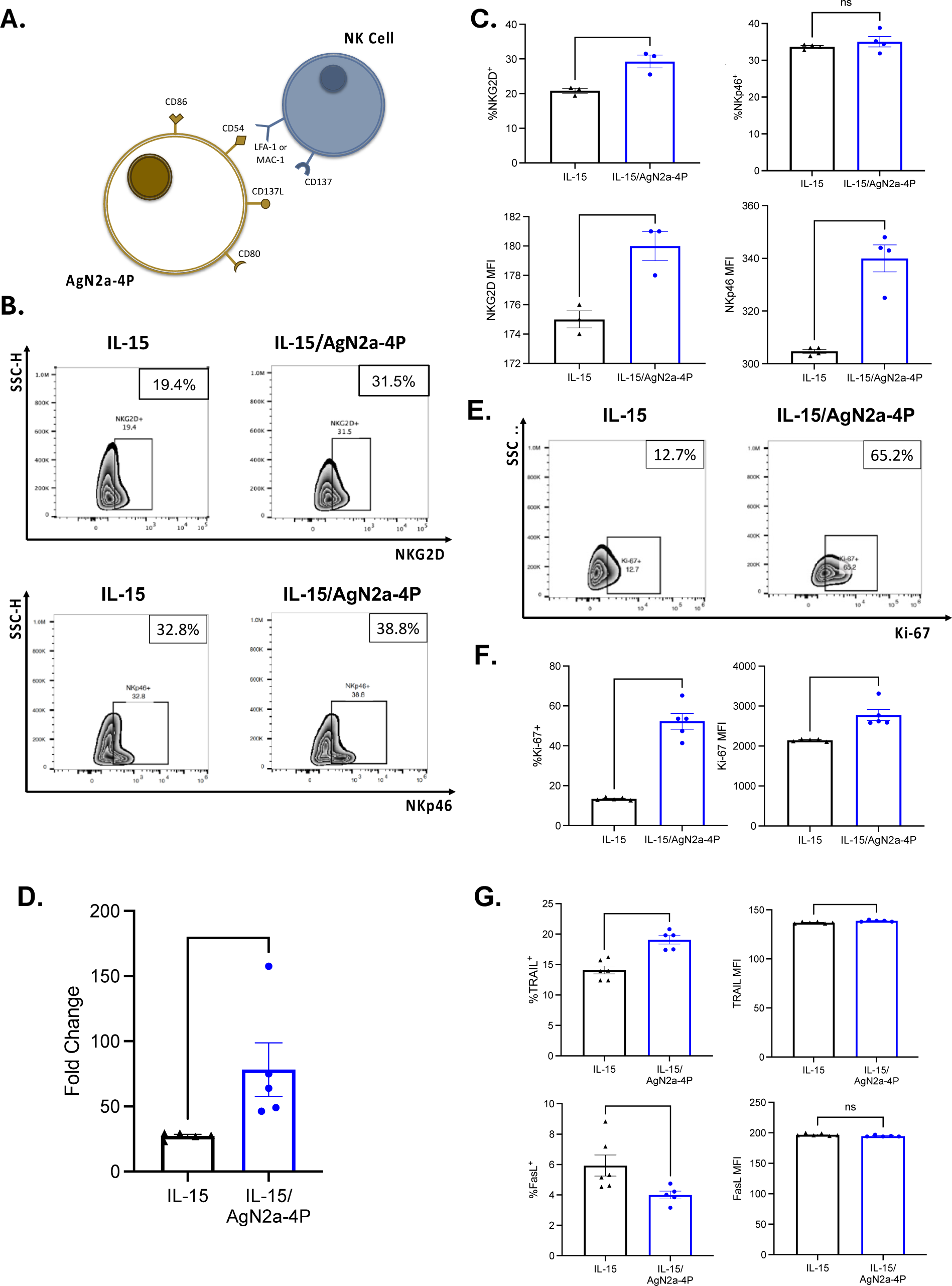
15-4P Stimulation increases NKG2D expression and murine NK cell proliferation compared to IL-15/IL-15Rα stimulation alone. (A) Schematic showing AgN2a-4P and NK cell receptor-ligand interactions. B6 NK cells are isolated and cultured with IL-15/IL-15Rα conjugate and with or without irradiated AgN2a-4P cells at a 1:1 ratio for 12 days. All expression markers were examined on CD3^-^NK1.1^+^ NK cells. (B) Dot plots of NKG2D (top) and NKp46 (bottom) on IL-15 NK cells (left) and 15-4P stimulated NK cells (right) are shown with percentage of gated cells. (C) Percent of NK cells positive for NKG2D (left) and NKp46 (right) and MFI below. (D) Fold change of IL-15 NK cells and 15-4P stimulated NK cell expansion at day 12 compared to day 0. (E) Dot plots of proliferation marker Ki-67 on IL-15 NK cells (left) and 15-4P stimulated NK cells (right) are shown with (F) percentage and MFI. (G) Percentage and MFI of TRAIL (top) and Fas-L (bottom) were assessed for each group. Data are representative of an experiment that was replicated two times. Representative dot plots examples of one replicate out of a minimum of 3 replicates. All bar graphs show individual experimental replicates plotted with mean and SEM (n = 3-5). Two-sided two-sample t tests were performed where indicated (* p < 0.05, ** p < 0.01, *** p < 0.001, ns = not significant).

To examine effector capacity, expansion with 15-4P resulted in a significantly higher percentage of TRAIL^+^ NK cells (19.06 ± 0.6823) compared to IL-15 NK cells (14.10 ± 0.6532) and a marked increase in TRAIL expression by MFI. Conversely, 15-4P stimulated NK cells exhibited a significant decrease in Fas-L^+^ cells (3.998 ± 0.2588) in contrast to IL-15 NK cells (5.935 ± 0.6996), and there was no difference in Fas-L expression by MFI (Fig. 1G). These findings indicate that exposure to AgN2a-4P during IL-15/IL-15Rα-based NK cell expansion results in increased expression of activation and proliferation markers compared to expansion with IL-15/IL-15Rα alone with a potential impact of cytotoxicity depending on if TRAIL versus FasL pathways are used to trigger apoptosis of NBL tumors.

The NBL TME is highly immunosuppressive, influencing NK cells through a complex interplay of activating and inhibitory receptor signaling. TIM-3, a marker of maturation in NK cells, interacts with its cognate ligands, including CEACAM-1, Gal-9, HMGB1, and PtdSer to exert inhibitory regulation in NK cells. We verified the surface expression of MHC I and Galectin-9 and CEACAM1 on murine NBL cell lines Neuro2a and NXS2. Neuro2a displayed a significantly higher percentage of CEACAM1+ cells (20.08% ± 0.9016) and Gal-9+ cells (95.10% ± 0.4235) compared to NXS2 (Fig. 2B). Both Neuro2a and NXS2 had low expression of MHC-I. The MFI of CEACAM1 and Galectin-9 was also notably higher on Neuro2a cells compared to NXS2 (Fig. 2C). Due to their intracellular expression, HMGB-1 and PtdSer mRNA were measured. Neuro2a showed significantly higher mRNA expression of both HMGB-1 and PtdSer compared to NXS2 (Supplementary Fig. 1A).

**Figure 2.**
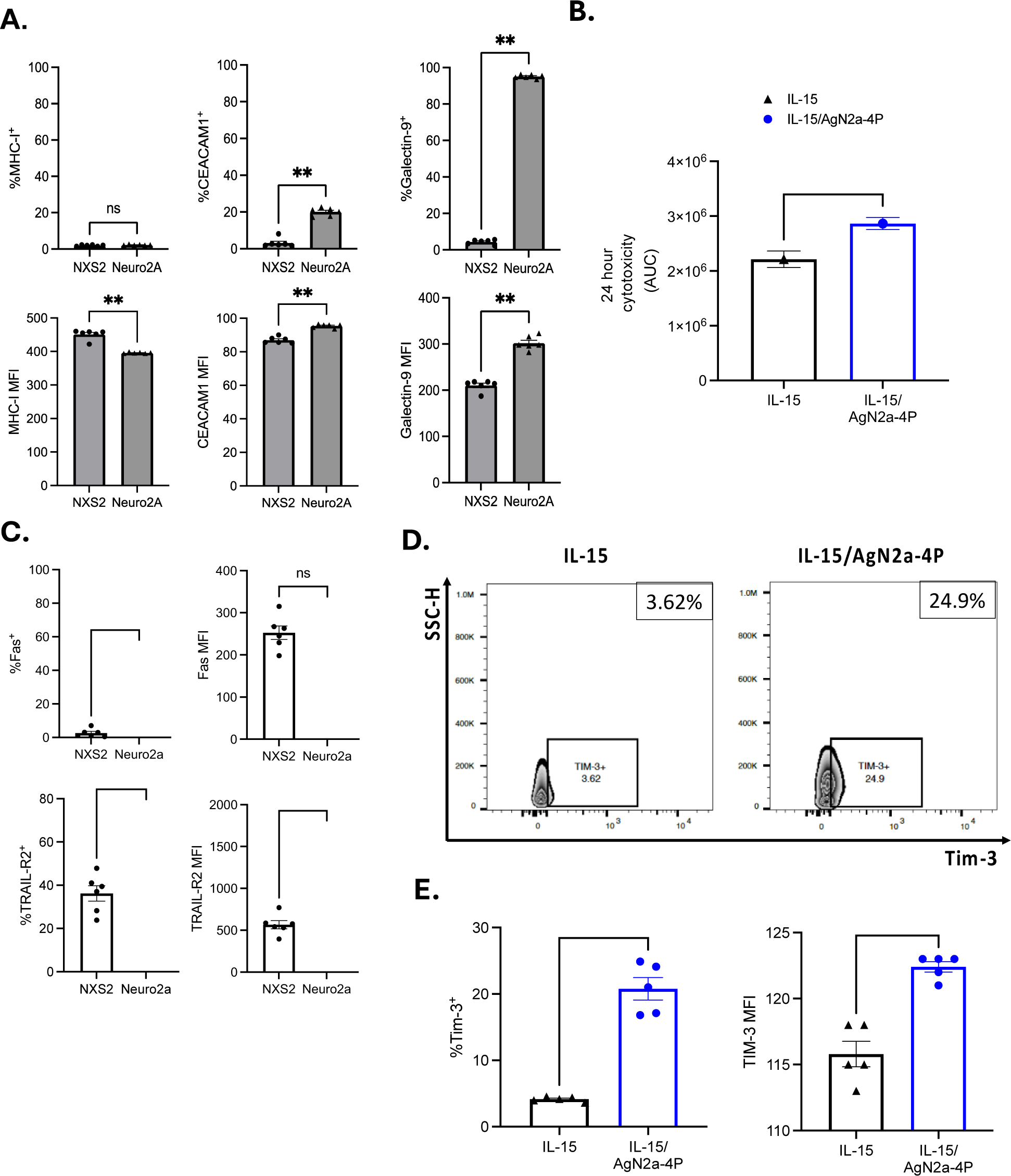
15-4P stimulated B6 NK cells show limited efficacy against neuroblastoma in vitro and increased expression of TIM-3. (A) NBL cell lines, NXS2 and Neuro2A, were interrogated for the presence of tumor ligands Ceacam-1, Galectin-9, MHC-I and with percentage (top) and MFI (bottom) shown for each marker. (B) IL-15 NK cells (black) or 15-4P stimulated NK cells (blue) were plated with neuroblastoma Neuro2A cells at an effector: target ratio of 10:1 with green fluorescent caspase 3/7-substrate for 24 hours. Total green object area (μm2/well) was measured by Incucyte real-time analysis and area under the curve (AUC) are shown for all groups. Mean with SEM shown for experimental replicates (n = 3 per group). (C) Fas (top) and TRAIL-R2 (bottom) percent positive cells and MFI are shown for NX2S and Neuro2a. (D) Representative dot plots for subsets of Tim-3^+^ IL-15 NK cells (left) and 15-4P stimulated NK cells (right) are shown. (E) Percentage (top), and MFI (bottom) are shown for the presence of Tim-3^+^ on IL-15 NK cells (left) and 15-4P stimulated NK cells (right). All the data are representative of an experiment that was replicated two times. Mean with SEM shown for experimental replicates (n = 6 per group). Two -sided two-sample t tests were performed (* p < 0.05, ** p < 0.01, ns = not significant).

Next, we evaluated the efficacy of 15-4P-stimulated ex-vivo activated allogeneic B6 NK cells (H-2b) against Neuro2a (H-2a) murine NBL to evaluate the impact of Ly49 mismatch on cytotoxicity. Neuro2a NBL targets were cocultured with either IL-15/IL-15Rα expanded allogeneic NK cells or 15-4P expanded allogeneic NK cells for 24 hours. Apoptotic Neuro2a cells were selected by size and fluorescence intensity of caspase 3/7 and quantified using live-cell imaging by the IncuCyte system. 15-4P-expanded allogeneic NK cells exhibited slightly increased target killing compared to IL-15-expanded allogeneic NK cells, despite showing a significant decrease in Fas-L expression (Fig. 2A).

Surface TIM-3 upregulation was confirmed on IL-15/IL-15Rα NK cells and 15-4P stimulated NK cells. Results showed a significantly higher percentage of TIM-3^+^ cells (20.78% ± 1.695) and TIM-3 expression by MFI in 15-4P stimulated NK cells relative to IL-15/IL-15Rα NK cells (4.134% ± 0.1592) (Fig. 2D-E). The elevated expression of TIM-3 on AgN2a-4P NK cells combined with elevated expression of putative TIM-3 ligands on NBL cells suggests this immune checkpoint may be a relevant axis governing NK cell function.

### 15-4P stimulated NK cells show enhanced cytotoxicity against NBL in vitro after TIM-3 immune checkpoint blockade

Initially, we measured IFNγ and TNFα mRNA levels to determine the impact of TIM-3 blockade on NK cell cytokine production after ex vivo expansion. We co-cultured IL-15 B6 allogeneic NK cells or 15-4P stimulated B6 allogeneic NK cells with isotype control or anti-TIM-3 antibody in the presence of Neuro2a NBL cells at an effector-to-target (E:T) ratio of 10:1 and assessed cytokine production. TNFα production was significantly increased in the 15-4P NK cells compared to IL-15 NK cells, however there was no increase after 15-4P NK cell with TIM-3 blockade after exposure to Neuro2a NBL. Similarly, we observed a notable increase in IFNγ production in 15-4P stimulated NK cells compared to IL-15 NK cells, but 15-4P NK cells showed no further increase in IFNγ (Supplementary Fig. 1B). Then, to investigate whether blocking TIM-3 could enhance the killing potential of 15-4P stimulated NK cells, we co-cultured 15-4P stimulated NK cells with Neuro2a or NXS2 NBL at an E:T ratio of 10:1 with isotype control or anti-TIM-3 and then assessed surface expression of activating receptors and death ligands as well as cytotoxicity. There was a high percentage of NKG2D+ and NKp46+ cells and elevated surface expression of NKG2D and NKp46 by MFI among 15-4P stimulated NK cells as seen in prior experiments, but these subsets were not enriched by TIM-3 blockade, suggesting NK cells were already maximally activated. Interestingly, in the presence of NXS2, 15-4P NK cells treated with TIM-3 blockade also show no difference in NKG2D and NKp46 expression by MFI as compared to 15-4P NK cells (Supplementary Fig. 2A). 15-4P NK cells exhibited a high percentage of TRAIL+ cells in the presence of the Neuro2a NBL, but the addition of anti-TIM-3 further augmented TRAIL expression. Moreover, 15-4P stimulated NK cells demonstrated significantly higher expression of TRAIL by MFI after TIM-3 blockade. Compared to 15-4P cells alone, TIM-3 blockade significantly enhanced both the percentage of Fas-L+ 15-4P NK cells, and Fas-L expression compared to 15-4P NK cells after Neuro2a exposure (Fig. 3A). In the presence of NXS2 we found no significant increase in TRAIL, Fas-L, granzyme B or perforin percent or expression, suggesting that blockade of exposure to TIM-3 ligands on NBL is required to augment allogeneic NK cell activation (Supplementary Fig. 2B).

**Figure 3.**
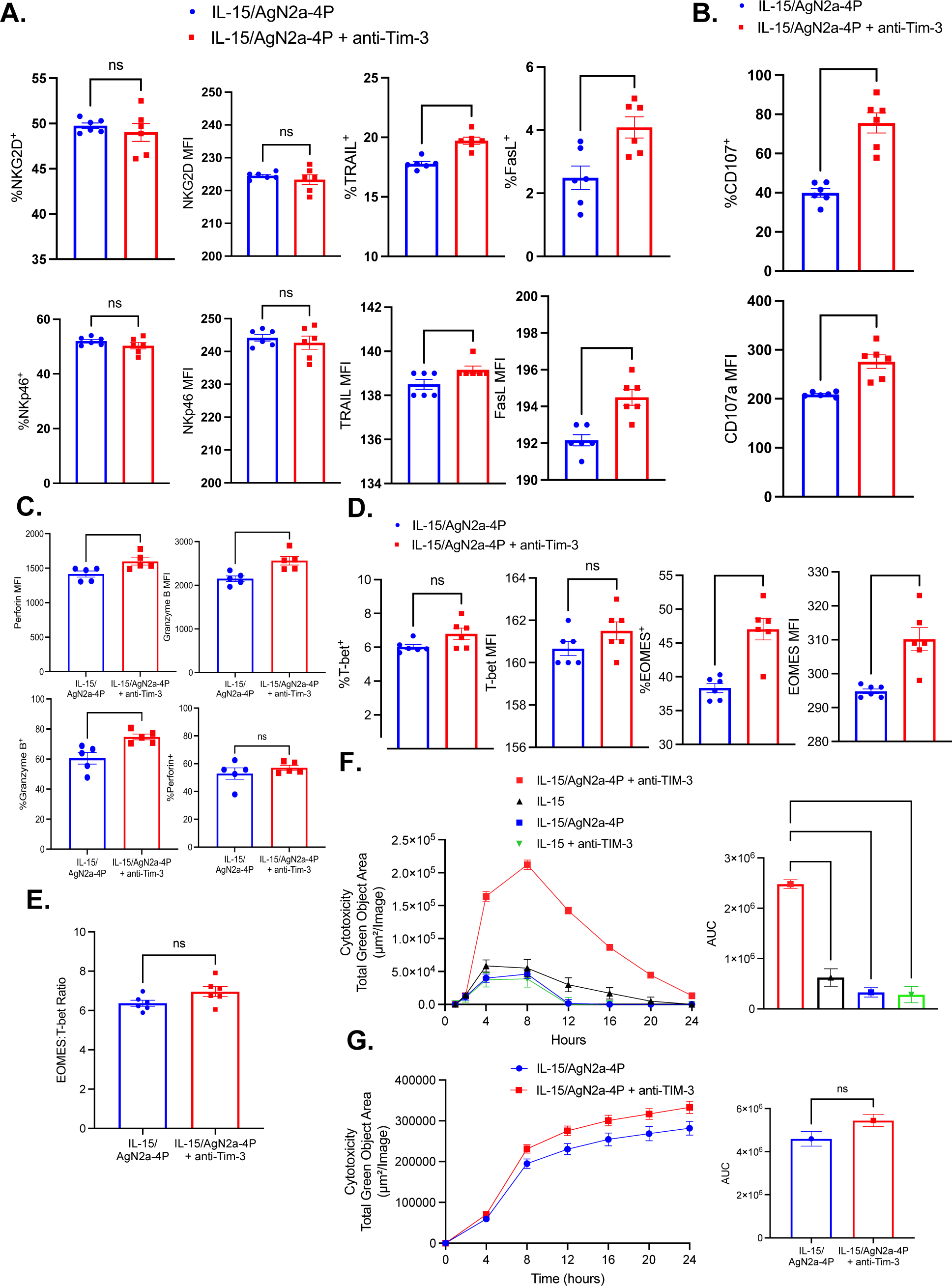
TIM-3 blockade enhances 15-4P stimulated NK cells cytotoxicity against NBL cells expressing TIM-3 ligands. 15-4P stimulated NK cells with isotype control (blue) or anti-TIM-3 blockade (red) were plated with Neuro2a NBL at an E:T ratio of 10:1 for 4 hours. NK cells were collected and analyzed by FACs. (A) NKG2D, NKp46, TRAIL, and Fas-L percentage positive (top) and MFI (top) for NK cells are shown. (B) NK cell degranulation was measured by CD107a percentage positive (top) and MFI (bottom) for 15-4P stimulated NK cells (blue) or 15-4P stimulated NK cells with anti-TIM-3 blockade (red). (C) Plots comparing percentage and MFI of granzyme B and perforin for 15-4P stimulated NK cells in the presence or absence of anti-TIM-3 blockade or IgG2a isotype control are shown. (D) Percentage and MFI of EOMES and T-bet are shown for all groups as well as (E) ratio of the percent of Eomes^+^ NK cells relative to the percent of T-bet^+^ NK cells. Mean with SEM shown for experimental replicates (n = 6). (F) For cytotoxicity experiments IL-15 NK cells with isotype control (black) or anti-Tim-3 blockade (green), and 15-4P stimulated NK cells with isotype control (blue) or TIM-3 blockade (red) were plated with Neuro2a NBL cells at an E:T ratio of 10:1 with green fluorescent caspase 3/7-substrate for 24 hours. (G) Cytotoxicity of 15-4P stimulated NK cells with isotype control or TIM-3 blockade against NXS2 NBL are shown. Cytotoxicity was determined by caspase 3/7 (total green object area (μm2/well)) tracking of tumor cell death with Incucyte real-time analysis, and area under the curve (AUC) was calculated for all groups. Two-sided two-sample t tests were performed for comparison between 2 groups. Comparisons between 3 or more groups was analyzed with Brown-Forsythe and Welch one-way ANOVA test with Dunnett’s T3 multiple comparisons test (* p < 0.05, ** p < 0.01, *** p < 0.001, **** p < 0.0001, ns = not significant).

Next, we evaluated cytotoxicity by quantifying CD107a degranulation on allogeneic NK cells following exposure to NBL. Interestingly, we found no discernible difference in degranulation between 15-4P stimulated NK cells with or without TIM-3 blockade in response to NXS2, which have lower expression of TIM-3 ligands (Supplementary Fig. 2C). However, TIM-3 blockade resulted in a significant increase in the percentage of CD107a+ 15-4P NK cells, as well as MFI of CD107a granules (Fig. 3B), in response to Neuro-2a. Additionally, we observed a significant increase in the percentage of granzyme B-positive cells following TIM-3 blockade, as well as a significant increase in both granzyme B and perforin expression (Fig. 3C).

To determine the impact of TIM-3 blockade on NK cell maturation, we assessed the expression of the transcription factors T-bet and Eomesodermin (Eomes). TIM-3 blockade did not impact T-bet percent or expression as 15-4P NK, and 15-4P NK with TIM-3 blockade, cells showed similarly low levels of T-bet. However, we observed a significant rise in both the percentage and expression of Eomes in 15-4P stimulated NK cells after exposure to TIM-3 blockade (47.05 ± 1.607) compared to 15-4P NK cells alone (38.33 ± 0.6756) (p = 0.0018). This was consistent with EOMES MFI (Fig. 3D). We determined the ratio of Eomes positive cells relative to T-bet in 15-4P stimulated NK cells and 15-4P stimulated NK cells with anti-TIM-3 after Neuro2a NBL exposure however there was no significant difference between the two groups (Fig 3E). Interestingly, TIM-3 blockade did not enhance EOMES expression in 15-4P NK cells after exposure to NXS2 NBL cells which have lower expression of TIM-3 ligands (Supplementary Fig. 2D). These findings suggest that TIM-3 blockade enriches for NK cells in early maturation [18].

Following treatment with anti-TIM-3, 15-4P stimulated NK cells exhibited heightened expression of death ligands and increased markers of degranulation, indicating a phenotype conducive to anti-tumor activity. Allogeneic NK cells were incubated with Neuro2a NBL with anti-TIM-3 or an isotype control and apoptotic NBL cells were quantified using Incucyte live cell imaging of caspase 3/7 expression. For all groups, peak killing occurred after the 8-hour mark. 15-4P NK cells treated with anti-TIM-3 antibody exhibited significantly higher killing compared to all other groups (Fig. 3F). The enhanced killing capacity of 15-4P NK cells was only observed in the presence of Neuro2a which has high expression of TIM-3 ligands, as we did not observe this effect against NXS2 (Fig. 3G).

### TIM-3 inhibition modulates gene expression in NK cells

We conducted a comparative evaluation of IL-15 allogeneic NK cells and 15-4P expanded allogeneic NK cells following exposure to the NBL TME to investigate whether TIM-3 blockade modulated NK cell gene expression. Total RNA was isolated from 15-4P stimulated NK cells NK cells co-cultured with Neuro2a in the presence of isotype controls or anti-TIM-3 for 4 hours. Using the NanoString/nCounter Mouse PanCancer Immune Profiling Panel, the expression of 760 genes were examined. The comparison between 15-4P stimulated NK cells and 15-4P stimulated NK cells treated with TIM-3 blockade showed 25 significantly upregulated genes, and 12 genes showed significant downregulation compared to 15-4P stimulated NK cells treated with an isotype control. Within the upregulated subset of genes, numerous genes associated with NK cell trafficking and target cell recognition (CCL4, CCL3, CCL1, XCL1, CXCR6), and the non-canonical NF-κB pathway (LTA, LTB, TNFSF10, TNFSF14, TNFSF8, RELB, TNF, TRAF3) [19] were observed. Both ICAM1, which encodes CD54, and TNFSF10, which encodes TRAIL, exhibited a fold change of 2.4 (with FDR p-values of 0.003 and 0.0024, respectively). Furthermore, genes encoding IL-15 and IL-21 receptors were upregulated, along with CD160, a NK cell-activating receptor that plays a significant role in modulating the cytokine response [20,21] Surprisingly, Prf1 (perforin) was downregulated in IL-15/AgN2a-4P/TIM-3 NK compared to 15-4P stimulated NK cells despite TRAIL upregulation (Fig. 4A-B; Supplementary Table 1). This alteration may suggest functional impairment due to exhaustion. Following the initial granule-mediated killing phase, NK cells may switch to death receptor signaling to sustain serial killing upon granule depletion [22,23]. NK cells use multiple cytotoxic pathways in tumor surveillance, transitioning between granule release and death receptor-mediated killing to initiate and sustain long-term cytotoxic activity. To assess the impact of multiple NK cell cytotoxicity pathways on the killing efficiency of 15-4P stimulated NK cells in the presence of TIM-3 blockade, we exposed allogeneic NK cells to Neuro2a NBL in the presence of blocking antibodies targeting NKG2D, TRAIL, and Fas-L. The inhibition of all three targets resulted in a significant reduction in NK cell killing. Notably, the blockade of Fas-L led completely ameliorated NK cell cytotoxicity against NBL compared to NKG2D or TRAIL blockade, suggesting there is a hierarchy of pathways used to mediate NK cytotoxicity of NBL when TIM-3 is engaged. (Fig. 4C). These data underscore the potential of TIM-3 blockade to enhance NK cell cytotoxicity against NBL, particularly in tumors with high expression of TIM-3 ligands, emphasizing the importance of considering the makeup of the NBL TME in therapeutic strategies.

**Figure 4.**
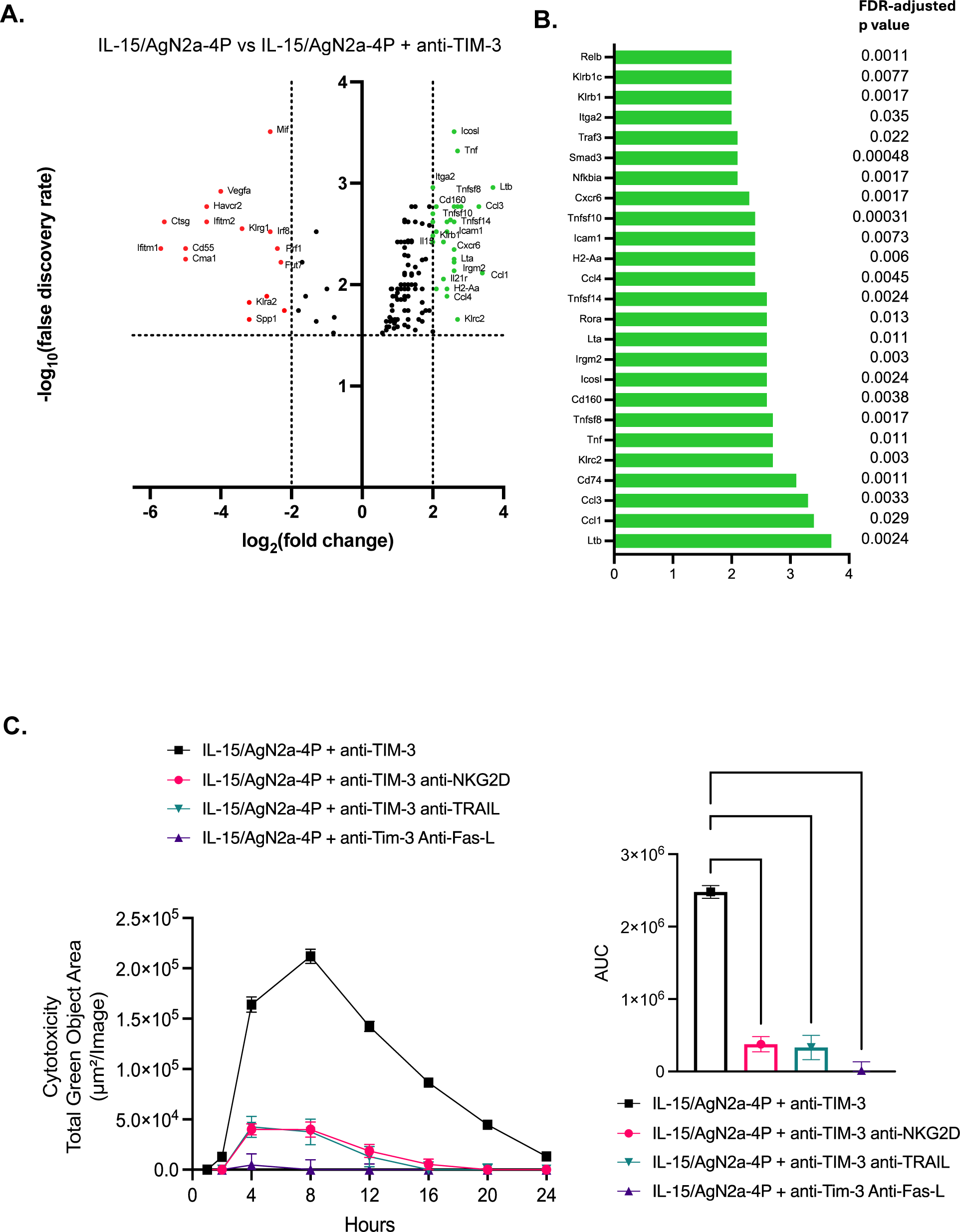
15-4P stimulated NK cells treated with anti-TIM-3 blockade respond against NBL through multiple cytotoxicity pathways. (A) Volcano plots representing the -log10 false discovery rate (FDR)-adjusted p-values and log2 fold changes for differentially expressed genes within the comparison of 15-4P NK cells with isotype control vs 15-4P NK cells with anti-TIM-3 antibody (right) (n = 2). The significance threshold for an FDR of ≤ 0.05 is represented by the horizontal line. Vertical lines represent genes with absolute fold change of ≥ 2. Upregulated genes are shown as green dots and downregulated genes are shown as red dots. (B) Upregulated DEGs from the comparison of 15-4P NK cells with isotype control vs 15-4P NK cells with anti-TIM-3 antibody, exhibiting an absolute fold change of ≥ 2, are plotted with respective fold change and p values. (C) 15-4P stimulated NK cells with anti-TIM-3 blockade (green) and anti-NKG2D blockade, anti-Fas-L blockade, or anti-TRAIL blockade were plated with Neuro2a NBL cells at an E:T ratio of 10:1 with green fluorescent caspase 3/7 - substrate for 24 hours. Cytotoxicity and AUC (D) are shown for all groups. All the data are representative of an experiment that was replicated two to three times. Mean with SEM shown for technical replicates (n = 5 per group). Comparisons between 3 or more groups was analyzed with Brown-Forsythe and Welch one-way ANOVA test with Dunnett’s T3 multiple comparisons test (**** p < 0.0001).

### Combination therapy of 15-4P stimulated allogeneic NK cells with anti-TIM-3 reduces NBL progression after allo-HCT

Given our observations regarding the positive impact of TIM-3 blockade on the phenotype and function of 15-4P stimulated NK cells in vitro, we hypothesized that this therapy could yield therapeutic benefits in vivo. However, the intricate pro-inflammatory environment post allo-HCT and the immunosuppressive NBL TME present additional challenges that could impede NK cell cytotoxic efficacy. To address this concern, we evaluated the potency of the combination allogeneic NK cell treatment against Neuro2a NBL following allo-HCT. Initially, we established an MHC-mismatched haploidentical allo-HCT model wherein lethally irradiated B6AJF1 (H-2b x H-2a) recipients received 5×10^6^ T cell-depleted B6 (H-2b) bone marrow with 1×10^4^ B6 T cell add back on day +0 to simulate residual T cells present in the donor graft during clinical T cell depletion protocols (e.g. CD3 or αβ-TCR depletion). On day +7, mice were subcutaneously inoculated with Neuro2a (H-2a) cells to mimic relapse post-HCT. Recipients then were infused with three weekly infusions of B6 IL-15 allogeneic NK cells or B6 15-4P stimulated allogeneic NK cells administered with either an isotype control or anti-TIM-3 treatment every 7 days starting on day +9 (Fig. 5A). Although TIM-3 blockade has not been tested after haploidentical allo-HCT, despite the presence of residual T cells in the donor bone marrow graft, we did not observe significant differences in GVHD-associated weight loss (Fig. 5B). Treatment with 15-4P allogeneic NK cells combined with TIM-3 blockade significantly prolonged overall survival compared to mice treated with 15-4P stimulated NK cells and an isotype control, IL-15 NK cells alone, or no treatment groups (Fig. 5C). By day 35, mice treated with 15-4P NK cells and anti-TIM-3 exhibited the most substantial reduction in tumor growth compared to those treated with an isotype control or NK cells alone (Fig. 5D). It is noteworthy that we did not observe differences between the 15-4P NK-treated group and the IL-15 NK-treated group, despite the disparities observed in vitro. This could be attributed to a combination of potential off-target effects of TIM-3 blockade and the presence of immunosuppressive factors within the TME, which may be being mitigated by TIM-3 blockade.

**Figure 5.**
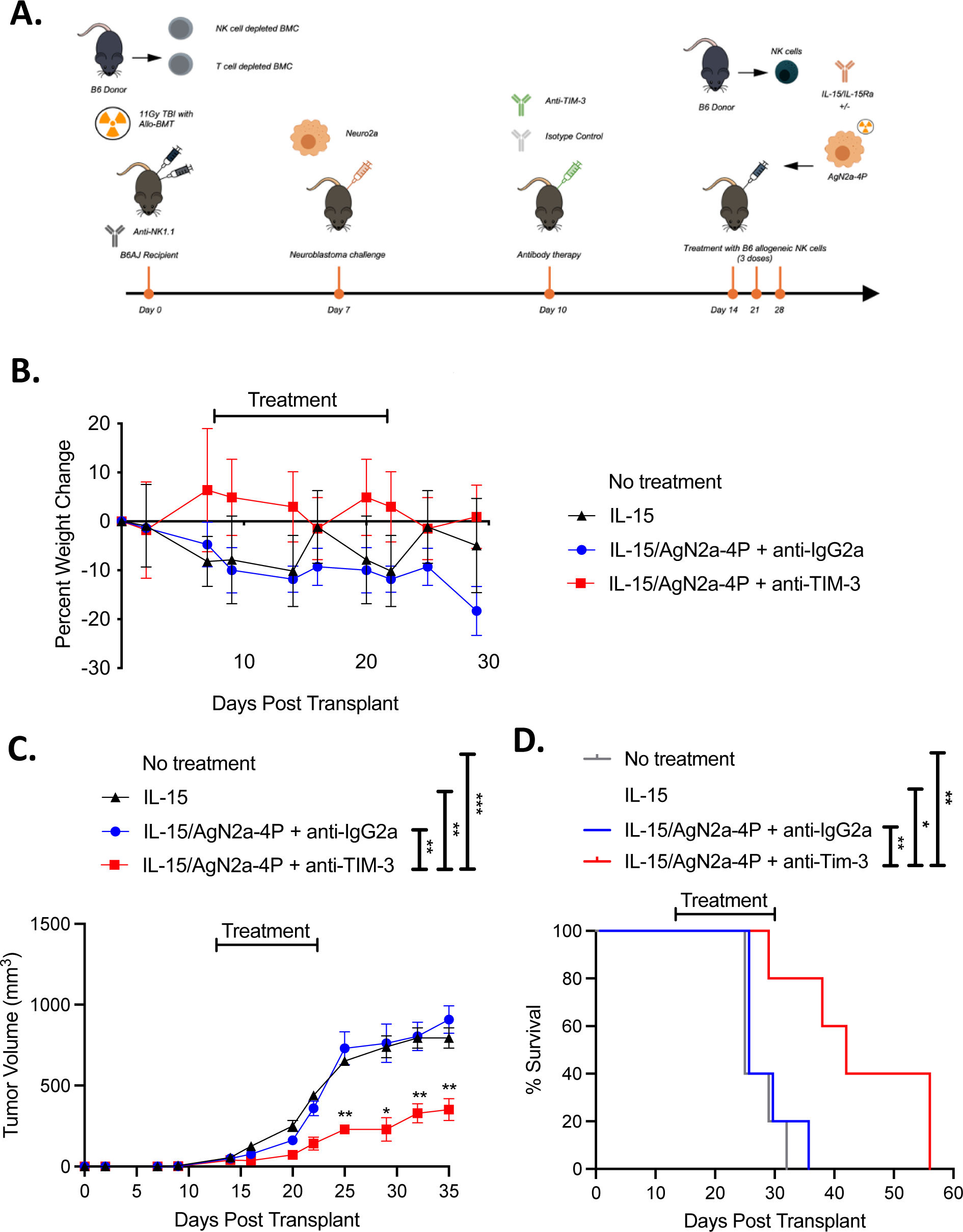
TIM-3 blockade with 15-4P stimulated allogeneic NK cells increase GVT without inducing GVHD after allo-HCT. (A) Schematic shown for post allo-HCT treatment of NBL relapse. (B) Percent weight change for mice treated with alloBMT and IL-15 B6 NK cells or 15-4P stimulated B6 NK cells with isotype or TIM-3 blockade are shown (n = 5 mice per group). (C) Overall survival for mice receiving alloBMT and no further treatment (NT) or treatment with 3 infusions of IL-15 B6 NK cells or 15-4P stimulated B6 NK cells with isotype or TIM-3 blockade are shown (n = 5 mice per group). (D) Tumor volume shown for with alloBMT and IL-15 B6 NK cells or 15-4P stimulated B6 NK cells with isotype or TIM-3 blockade are shown (n = 5 mice per group). All the data are representative of an experiment that was replicated two times. Tumor volume was calculated by as V = ½ (Length × Width^2^). Bars and points with error bars show mean with SEM. One-way ANOVA and Kruskal-Wallis with Dunn’s multiple comparisons tests were performed (* p < 0.05, ** p < 0.01, *** p < 0.001, ns = not significant).

Given the known role of TIM-3 blockade in promoting T cell responses and the presence of a low dose of T cells in the graft, we sought to delineate the contributions of NK cells and T cells to GVT effects and overall survival in our allo-HCT model by depletion of these cell subsets prior to treatment. In our T cell depletion studies, B6AJF1 mice received T-depleted B6 bone marrow without T cell add-back. In NK depletion studies, B6AJF1 mice received NK-depleted B6 bone marrow with T cell add-back. Since NK cells typically are the first lymphocyte to reconstitute after allo-HCT, anti-NK1.1 antibody was administered intraperitoneally every 4 days to maintain NK cell depletion. Subsequently, mice were inoculated with Neuro2A cells and treated with anti-TIM-3 antibody alone or B6 15-4P stimulated allogeneic NK cells with anti-TIM-3 antibody (Fig. 6A). Consistently, there were no noticeable differences concerning GVHD-associated weight loss (Fig. 6B). NK cell depletion led to a significant decrease in overall survival, while T cell-depleted mice exhibited significantly increased survival like that of mice treated with 15-4P stimulated NK cells and TIM-3 blockade (Fig. 6C). Furthermore, NK cell depletion resulted in larger tumors in all mice, with no significant difference compared to the control, whereas T cell depletion led to a significant decrease in tumor growth (Fig. 6D). These findings suggest that while T cells are present in the graft, NK cells may play a more substantial role in the survival benefit conferred by 15-4P stimulated NK cells, possibly due to the limited number of T cells compared to NK cells resulting from repeated adoptive transfers. Moreover, these results underscore the crucial role of NK cells as the primary drivers of TIM-3-induced therapeutic benefits against NBL after allo-HCT, given that similar outcomes were not observed with anti-TIM-3 treatment alone (Supplementary Fig. 3A-B). Nonetheless, our approach facilitates potent GVT effects against NBL without exacerbating GVHD, through the combination therapy of anti-TIM-3 with adoptive transfer of ex vivo expanded allogeneic NK cells.

**Figure 6.**
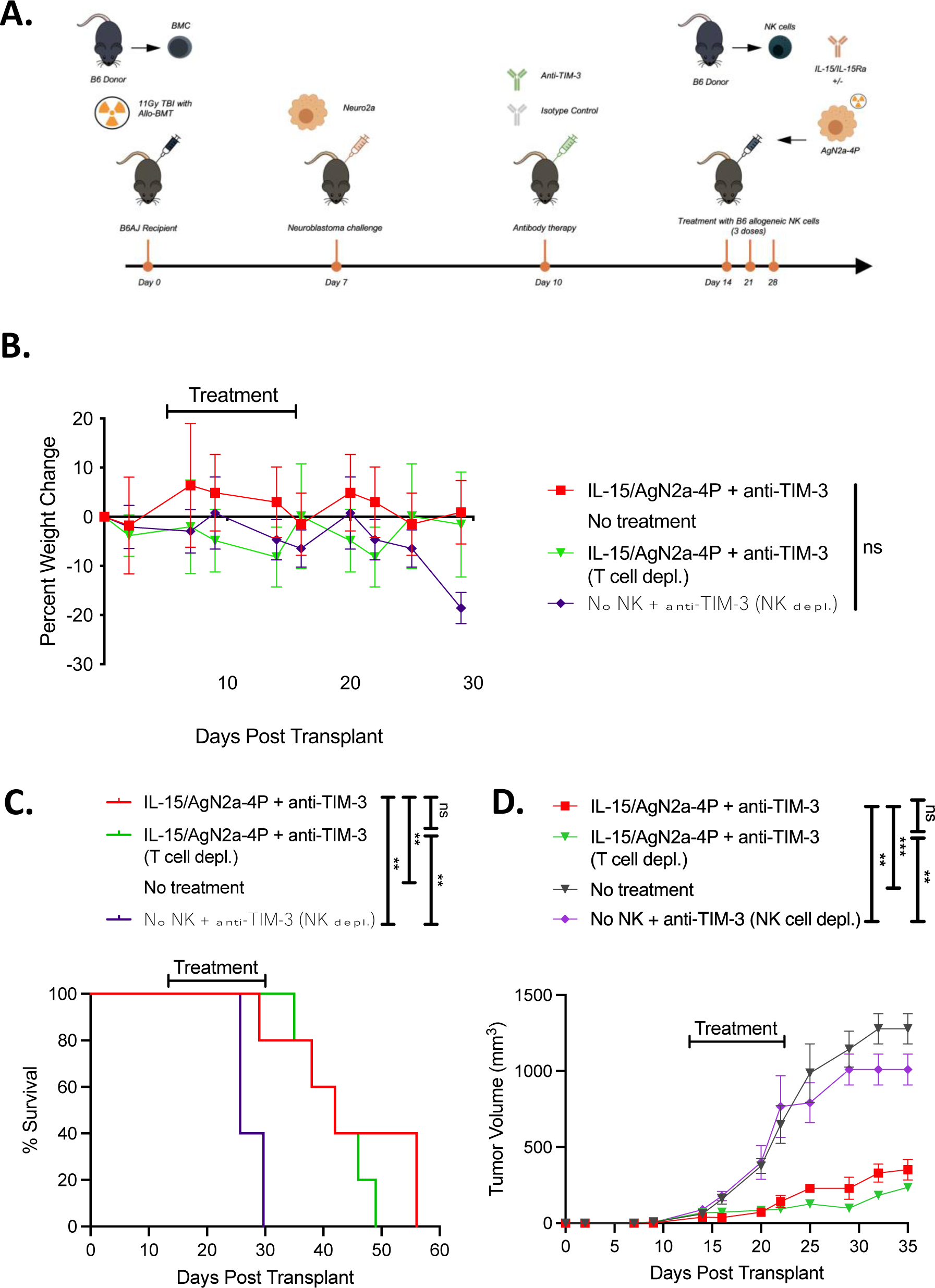
NK cells are necessary for the GVT effect mediated by TIM-3 blockade against NBL relapse after allo-HCT. (A) Schematic shown for post alloBMT treatment of NBL relapse with NK and T cell depletion. (B) Percent weight change, (C) overall survival and (D) tumor volume is shown for mice receiving alloBMT with no further treatment or treatment with 3 infusions of 15-4P stimulated B6 NK and TIM-3 blockade, T cell depleted alloBMT with 3 infusions of 15-4P stimulated B6 NK and TIM-3 blockade, or NK cell depleted alloBMT with TIM-3 blockade. (n = 5 mice per group). Tumor volume was calculated by as V = ½ (Length × Width^2^). Bars and points with error bars show mean with SEM. one-way ANOVA and Kruskal-Wallis with Dunn’s multiple comparisons tests were performed (** p < 0.01, *** p < 0.001, ns = not significant).

## DISCUSSION

T cell depleted (TCD) allo-HCT has recently gained traction as a safe alternative to standard unmanipulated allo-HCT in order to leverage the GVT effect mediated by donor-derived innate effector cells, like NK cells, while mitigating lethal GVHD [24]. While graft manipulation techniques like CD3 or αβ-TCR depletion have achieved up to 5 Log_10_ depletion of T cells, there are always depletion resistant CD8+ or CD4+ T-cells to consider toward impacting GVT or GVHD [25]. A major limitation of TCD allo-HCT is the increased risk of infectious complications and relapse due to delayed immune reconstitution of T cells. Preclinically, we have previously explored the use of T cell-depleted haploidentical allo-HCT in combination with adoptively transferred ex vivo activated NK cells and an anti-GD2 immunocytokine[18]. In this study we found that 15-4P ex vivo expanded NK cells have increased potency against NBL target cells, especially when combined with TIM-3 blockade. CD54 and CD137L co-stimulation have never been combined to activate murine NK cells ex vivo. While CD54+ CD137L+ AgN2a-4P cells were initially designed as an in vivo vaccine for stimulating T cells, we demonstrate for the first time that combining IL-15/IL-15Ra with AgN2a-4P exposure ex vivo significantly boosts NK cell proliferation and activation, allowing for adoptive transfer of highly potent immune effector cells in the allo-HCT setting.

AgN2a-4P stimulation increased both NKG2D and TRAIL expression on NK cells. 15-4P-stimulated NK cells exhibited increased IFNv production compared to IL-15/IL-15RA stimulation alone. Prior studies done in a syngeneic model by Jing et al. demonstrated that AgN2a-4P vaccination enacted an anti-NBL response through CD8^+^ and CD4+ T cells mediated response, despite their tumor target being the parental line AgN2a, which does not express MHC class II [26]. While studies have shown that CD4^+^ T cells can kill tumor targets through an indirect mechanism involving IFN-gamma directed elimination, the contribution of NK cells to the anti-NBL response cannot be ruled out [27]. NK cell reconstitution occurs rapidly following HSCT, and it is possible that AgN2a-4P vaccination induced an NK cell mediated anti-tumor response and increased IFNγ production which indirectly facilitated CD8+ T cell and macrophage anti-tumor responses [26]. Additionally, we found that 15-4P stimulation significantly enhanced NK cell mediated lysis of Neuro2a NBL tumor compared to IL-15 NK cells in vitro. IL-15 exposure results in an upregulation of TIM-3 expression where TIM-3 can act as a negative regulator of activation to limit autoreactivity in innate immune cells and maintain self-tolerance [28,29]. Although increased TIM-3 upregulation is associated with activated NK cells, cytotoxicity is significantly impaired upon crosslinking of TIM-3 with cognate ligands [15]. In addition to TIM-3 regulation of NK cell activation, inhibitory signals in the NBL microenvironment can dampen the NK cell response through mechanisms including blockade of target cell adhesion and inhibition of LFA-1 activation subjugate a weaker activating stimulus, tilting the NK cell response towards inactivation [30,31]. TIM-3 ligands such as galectin-9, CEACAM-1, HMGB1, and PtdSer are expressed at different levels on and within NBL cells and are present in the TME as they are ancillary to pathogenesis [32–34]. Our results showed that multiple TIM-3 ligands were expressed at a higher proportion on Neuro2a, but not NXS2, suggesting that TIM-3 ligand exposure is cell line dependent and may only be upregulated in a subset of NBL patients. However, exposure to inhibitory signals from TIM-3 ligands can originate from tumor associated macrophages as well as other immune cell types within the microenvironment [35].

To better understand the mechanisms driving 15-4P NK cells treated with TIM-3 blockade against NBL, we analyzed upregulation of 25 genes and downregulation of 12 genes in 15-4P stimulated NK cells compared to 15-4P stimulated NK cells alone, including ICAM1 and TNFSF10 encoding CD54 and TRAIL, respectively. Within the upregulated subset of genes, numerous genes were found to be associated with NK cell trafficking and target cell recognition (CCL4, CCL3, CCL1, XCL1, CXCR6) in 15-4P stimulated NK cells with TIM-3 blockade compared 15-4P NK cells alone. Modulating chemokine receptors may contribute to increased NK cell motility, thereby allowing 15-4P NK cells enhanced ability to track Neuro2a NBL cells in the absence of TIM-3 inhibition [36]. Interestingly, we saw increased expression of multiple genes involved in the TNFR2 non-canonical NF-kB pathway (LTA, LTB, TNFSF10, TNFSF14, TNFSF8, RELB, TNF, TRAF3) and in DAP12 signaling (H2-Aa, KLRC2) which suggests the involvement of NKG2D activation in anti-TIM-3 15-4P NK cell mediated killing. Functional cytotoxicity studies confirmed the ability of 15-4P NK cells treated with TIM-3 blockade to kill Neuro2A targets, which were reversed when blocking NKG2D, TRAIL, or Fas-L. Interestingly, we observed a downregulation of Prf1 (perforin) in 15-4P NK cells treated with TIM-3 blockade compared to 15-4P stimulated NK cells, while TNFSF10 (TRAIL) showed upregulation. This may suggest functional impairment in 15-4P NK cells treated with TIM-3 blockade, potentially due to exhaustion. However, the upregulation of TRAIL and the non-canonical NF-κB pathway intermediates suggests a compensatory mechanism, where NK cells engage the death receptor pathway for killing upon granule depletion, thereby sustaining serial killing of target cells [22].

Several studies have indicated that blocking TIM-3 interactions during hepatitis B virus (HBV) infection, as well as in melanoma, AML, and multiple myeloma resulted in enhanced NK cell cytotoxicity [37–40]. Conversely, other studies have reported enhanced IFN-γ production without a corresponding improvement in NK cell degranulation suggesting that TIM-3 engagement can shift the NK cell response between positive and negative signaling events through differential tyrosine phosphorylation of the cytoplasmic tail [13,41,42]. We observed increased IFN-γ production in 15-4P stimulated NK cells, which indicated that exposure to AgN2a-4P during ex vivo expansion shifts NK cells to a more immature phenotype that is still capable of robust cytotoxicity. However, the addition of TIM-3 blockade did not further enhance IFN-γ and TNF-α production in 15-4P NK cells. TIM-3 receptor-ligand engagement has been proposed to inhibit the ERK and NFκB pathways in NK cells, yet NFAT signaling remains unaffected, thereby enabling NK cells to sustain cytokine production [43]. Nevertheless, further investigation is warranted to fully elucidate the underlying mechanism behind the effect of TIM-3 on cytokine production in NK cells.

We validated our observations in a haploidentical allo-HSCT murine model of NBL relapse to establish pre-clinically the combination of adoptive transfer of NK cells and TIM-3 blockade treatment. We observed that the combination of NK cell donor lymphocyte infusions and TIM-3 blockade treatment as a viable therapeutic platform. Mice treated with multiple infusions of 15-4P stimulated NK cells and anti-TIM-3 antibody showed significantly smaller NBL tumors and prolonged survival. However, tumors began to grow shortly after treatment ceased. Prior studies show that following IL-15 expansion, allogeneic ex-vivo expanded NK cells remain detectable for up to 16 days after infusion in a murine model [44]. The limited persistence of adoptive NK cell therapies may require frequent infusions to effectively impact clinical outcomes, highlighting a limitation of this therapeutic approach. However, clinical studies have demonstrated favorable outcomes with the use of NK cell infusions combined with immunotherapy as an effective bridge therapy before consolidation [45]. Future studies may aim to generate a sustained NK cell response by designing an in vivo tumor vaccine with costimulatory molecules and/or cytokines specific to activating NK cells.

TIM-3 also plays a crucial role in modulating immune responses during acute GVHD following allogeneic hematopoietic stem cell transplantation (allo-HSCT). Donor T cells in the allogeneic setting rapidly upregulate TIM-3 expression. Inhibition of the TIM-3/Gal-9 interaction has been shown to lead to increased proliferation of T cells and heightened GVHD lethality. However, TIM-3 inhibition after regulatory T cell (Treg) depletion has been shown to increase IFN-γ production and reduce GVHD severity [46]. In our experimental model, mice received T-depleted BMCs with 1×10^3^ T cells added back to the graft. Notably, GVHD development was absent despite the use of TIM-3 blockade in the presence of residual donor T cells in the bone marrow graft. We then investigated whether the anti-NBL response was driven by NK cells or T cells in the graft. Only NK cell depletion reversed the clinical benefit of TIM-3 blockade, leading to larger tumors and worse survival compared to T cell depletion, which had no effect. It is possible that the 15-4P NK cell infusions increased IFNγ production while mediating GVT responses, which indirectly facilitated CD8+ T cell and macrophage anti-tumor responses as well [26]. Nevertheless, these results support the conclusion that the anti-NBL response is primarily driven by NK cells, although the potential contribution of blocking TIM-3+ macrophages and DCs on NBL clearance needs to be explored.

In summary, this study explored the combination of ex vivo activated NK cells with TCD allo-HCT and revealed the enhanced NK cell potency against relapsed NBL, particularly when paired with TIM-3 blockade. TCD allo-HCT has emerged as a safe and effective treatment for multiple hematologic malignancies, and in combination with cell-based immunotherapy may become a platform for treating solid tumors to maximize the GVT effect without exacerbating lethal GVHD. In a post-allo HCT relapsed NBL murine model, the combination of adoptive NK therapy with TIM3 blockade attenuated tumor growth and prolonged survival, without inducing GVHD, emphasizing the pivotal role of NK cells in mediating anti-NBL responses. Further studies are warranted to test the benefits of this combination therapy in a humanized mouse model against patient derived NBL samples and in combination with other emerging immune checkpoint inhibitors.

## List of Abbreviations

alloBMT: allogeneic bone marrow transplant
DEG: differentially expressed gene
E: T: effector: target
FDR: false discovery rate
GVHD: graft-versus-host disease
GVT: graft-versus-tumor
HCT: hematopoietic cell transplant
IFNγ: interferon gamma
KIR: killer-immunoglobulin-like receptor
MFI: median fluorescence intensity
MHC: major histocompatibility complex
NBL: neuroblastoma
NK: natural killer
TCD: T cell depleted
TME: tumor microenvironment
TNF: tumor necrosis factor

## Availability of data and material

The datasets used and/or analyzed during the current study are available from the corresponding author on reasonable request.

## Authors’ contributions

AEQ was responsible for experimental design, execution, and analysis of data, and drafted the manuscript. LSh isolated RNA and performed quality control for microarray analysis. NRM, CAK, AAG, JMK, MMC, SPR, KET, PDB, LSo and NJH contributed to experimental design and data collection. TJK and BDJ provided critical reagents and intellectual input. CMC provided experimental design, analysis of data, and edited the manuscript. All authors read and approved the final manuscript.

## Acknowledgements

We thank the Division of Hematology, Oncology and Bone Marrow Transplant in the Department of Pediatrics and Carbone Cancer Center at the University of Wisconsin-Madison for their ongoing support. We thank the University of Wisconsin Carbone Cancer Center Flow Cytometry Laboratory, supported by P30 CA014520, Small Animal Imaging and Radiotherapy Facility, and Translational Research Initiatives in Pathology, and Biomedical Research Model Services for use of their facilities and services.

## REFERENCES

1 Matthay KK, Maris JM, Schleiermacher G, et al. Neuroblastoma. Nat Rev Dis Primer. 2016;2:16078.

2 DuBois SG, Macy Margaret E, Henderson TO. High-Risk and Relapsed Neuroblastoma: Toward More Cures and Better Outcomes. Am Soc Clin Oncol Educ Book. 2022;768–80.

3 McNerney KO, Karageorgos SA, Hogarty MD, et al. Enhancing Neuroblastoma Immunotherapies by Engaging iNKT and NK Cells. Front Immunol. 2020;11:873.

4 Pritchard J, Cotterill SJ, Germond SM, et al. High dose melphalan in the treatment of advanced neuroblastoma: results of a randomised trial (ENSG-1) by the European Neuroblastoma Study Group. Pediatr Blood Cancer. 2005;44:348–57.

5 Berthold F, Boos J, Burdach S, et al. Myeloablative megatherapy with autologous stem-cell rescue versus oral maintenance chemotherapy as consolidation treatment in patients with high-risk neuroblastoma: a randomised controlled trial. Lancet Oncol. 2005;6:649–58.

6 Hale GA, Arora M, Ahn KW, et al. Allogeneic hematopoietic cell transplantation for neuroblastoma: the CIBMTR experience. Bone Marrow Transplant. 2013;48:1056–64. doi: 10.1038/bmt.2012.284

7 Federmann B, Bornhauser M, Meisner C, et al. Haploidentical allogeneic hematopoietic cell transplantation in adults using CD3/CD19 depletion and reduced intensity conditioning: a phase II study. Haematologica. 2012;97:1523–31.

8 Bethge WA, Faul C, Bornhäuser M, et al. Haploidentical allogeneic hematopoietic cell transplantation in adults using CD3/CD19 depletion and reduced intensity conditioning: An update. Blood Cells Mol Dis. 2008;40:13–9.

9 Blavier L, Yang R-M, DeClerck YA. The Tumor Microenvironment in Neuroblastoma: New Players, New Mechanisms of Interaction and New Perspectives. Cancers. 2020;12:2912.

10 Khalil M, Wang D, Hashemi E, et al. Implications of a ‘Third Signal’ in NK Cells. Cells. 2021;10:1955.

11 Meza Guzman LG, Keating N, Nicholson SE. Natural Killer Cells: Tumor Surveillance and Signaling. Cancers. 2020;12:952.

12 Olson JA, Leveson-Gower DB, Gill S, et al. NK cells mediate reduction of GVHD by inhibiting activated, alloreactive T cells while retaining GVT effects. Blood. 2010;115:4293– 301.

13 Gleason MK, Lenvik TR, McCullar V, et al. Tim-3 is an inducible human natural killer cell receptor that enhances interferon gamma production in response to galectin-9. Blood. 2012;119:3064–72.

14 So EC, Khaladj-Ghom A, Ji Y, et al. NK cell expression of Tim-3: *First impressions matter*. Immunobiology. 2019;224:362–70.

15 Ndhlovu LC, Lopez-Vergès S, Barbour JD, et al. Tim-3 marks human natural killer cell maturation and suppresses cell-mediated cytotoxicity. Blood. 2012;119:3734–43.

16 Navabi S sadat, Doroudchi M, Tashnizi AH, et al. Natural Killer Cell Functional Activity After 4-1BB Costimulation. Inflammation. 2015;38:1181–90.

17 Vahlne G, Becker S, Brodin P, et al. IFN-γ Production and Degranulation are Differentially Regulated in Response to Stimulation in Murine Natural Killer Cells. Scand J Immunol. 2008;67:1–11.

18 Lieberman NAP, DeGolier K, Haberthur K, et al. An Uncoupling of Canonical Phenotypic Markers and Functional Potency of Ex Vivo-Expanded Natural Killer Cells. Front Immunol. 2018;9:150.

19 Sun S-C. Non-canonical NF-κB signaling pathway. Cell Res. 2011;21:71–85.

20 Tu TC, Brown NK, Kim T-J, et al. CD160 is essential for NK-mediated IFN-γ production. J Exp Med. 2015;212:415–29.

21 Le Bouteiller P, Tabiasco J, Polgar B, et al. CD160: a unique activating NK cell receptor. Immunol Lett. 2011;138:93–6.

22 Prager I, Liesche C, van Ooijen H, et al. NK cells switch from granzyme B to death receptor–mediated cytotoxicity during serial killing. J Exp Med. 2019;216:2113–27.

23 Pipkin ME, Rao A, Lichtenheld MG. The transcriptional control of the perforin locus. Immunol Rev. 2010;235:55–72.

24 Flaadt T, Ladenstein RL, Ebinger M, et al. Anti-GD2 Antibody Dinutuximab Beta and Low-Dose Interleukin 2 After Haploidentical Stem-Cell Transplantation in Patients With Relapsed Neuroblastoma: A Multicenter, Phase I/II Trial. J Clin Oncol. 2023;41:3135–48.

25 Bueno V, Pestana JOM. The role of CD8+ T cells during allograft rejection. Braz J Med Biol Res Rev Bras Pesqui Medicas E Biol. 2002;35:1247–58.

26 Jing W, Orentas RJ, Johnson BD. Induction of Immunity to Neuroblastoma Early after Syngeneic Hematopoietic Stem Cell Transplantation Using a Novel Mouse Tumor Vaccine. Biol Blood Marrow Transplant J Am Soc Blood Marrow Transplant. 2007;13:277–92.

27 Mumberg D, Monach PA, Wanderling S, et al. CD4+ T cells eliminate MHC class II-negative cancer cells in vivo by indirect effects of IFN-γ. Proc Natl Acad Sci U S A. 1999;96:8633–8.

28 Han G, Chen G. Tim-3: An Activation Marker and Activation Limiter of Innate Immune Cells. Front Immunol. 2013;4. doi: 10.3389/fimmu.2013.00449

29 Khan M, Arooj S, Wang H. NK Cell-Based Immune Checkpoint Inhibition. Front Immunol. 2020;11.

30 Kumar S. Natural killer cell cytotoxicity and its regulation by inhibitory receptors. Immunology. 2018;154:383–93.

31 Das A, Long EO. Lytic Granule Polarization, rather than Degranulation, is the Preferred Target of Inhibitory Receptors in NK Cells. J Immunol Baltim Md 1950. 2010;185:10.4049/jimmunol.1001220.

32 Mari E, Zicari A, Fico F, et al. Action of HMGB1 on miR-221/222 cluster in neuroblastoma cell lines. Oncol Lett. 2016;12:2133–8.

33 Liu Y, Song L. HMGB1-induced autophagy in Schwann cells promotes neuroblastoma proliferation. Int J Clin Exp Pathol. 2015;8:504–10.

34 Poliani PL, Mitola S, Ravanini M, et al. CEACAM1/VEGF cross-talk during neuroblastic tumour differentiation. J Pathol. 2007;211:541–9.

35 Gao X, Zhou S, Qin Z, et al. Upregulation of HMGB1 in tumor-associated macrophages induced by tumor cell-derived lactate further promotes colorectal cancer progression. J Transl Med. 2023;21:53.

36 Krämer B, Schulte D, Körner C, et al. Regulation of NK cell trafficking by CD81. Eur J Immunol. 2009;39:3447–58.

37 Jiang W, Li F, Jiang Y, et al. Tim-3 Blockade Elicits Potent Anti-Multiple Myeloma Immunity of Natural Killer Cells. Front Oncol. 2022;12:739976.

38 Rakova J, Truxova I, Holicek P, et al. TIM-3 levels correlate with enhanced NK cell cytotoxicity and improved clinical outcome in AML patients. Oncoimmunology. ;10:1889822.

39 da Silva IP, Gallois A, Jimenez-Baranda S, et al. Reversal of NK-cell exhaustion in advanced melanoma by Tim-3 blockade. Cancer Immunol Res. 2014;2:410–22.

40 Ju Y, Hou N, Meng J, et al. T cell immunoglobulin- and mucin-domain-containing molecule-3 (Tim-3) mediates natural killer cell suppression in chronic hepatitis B. J Hepatol. 2010;52:322–9.

41 Anderson AC, Anderson DE, Bregoli L, et al. Promotion of tissue inflammation by the immune receptor Tim-3 expressed on innate immune cells. Science. 2007;318:1141–3.

42 Lee J, Su EW, Zhu C, et al. Phosphotyrosine-dependent coupling of Tim-3 to T-cell receptor signaling pathways. Mol Cell Biol. 2011;31:3963–74.

43. 43 Yu X, Lang B, Chen X, et al. The inhibitory receptor Tim-3 fails to suppress IFN-γ production via the NFAT pathway in NK-cell, unlike that in CD4+ T cells. BMC Immunol. 2021;22:25.

44 Cho MM, Song L, Quamine AE, et al. CD155 blockade enhances allogeneic natural killer cell-mediated antitumor response against osteosarcoma. BioRxiv Prepr Serv Biol. 2024;2023.06.07.544144.

45 Björklund AT, Carlsten M, Sohlberg E, et al. Complete Remission with Reduction of High-Risk Clones following Haploidentical NK-Cell Therapy against MDS and AML. Clin Cancer Res. 2018;24:1834–44.

46 Veenstra RG, Taylor PA, Zhou Q, et al. Contrasting acute graft-versus-host disease effects of Tim-3/galectin-9 pathway blockade dependent upon the presence of donor regulatory T cells. Blood. 2012;120:682–90.

