## Supplemental Figures for "TIM-3 blockade enhances ex vivo stimulated allogeneic NK cell therapy for relapsed murine neuroblastoma after hematopoietic cell transplant"

### SUPPLEMENTARY TABLES AND FIGURES

#### TIM-3 blockade enhances tumor vaccine-stimulated natural killer cell therapy for relapsed/refractory murine neuroblastoma after allogeneic transplant

Aicha E Quamine , Nicholas R Mohrdieck, Chloe A King, Anastasia A Griggs, Jillian M Kline, Monica M Cho, Lei Shi, Longzhen Song, Katharine E Tippins, Paul D Bates, Nicholas J Hess, Christian M Capitini

**Supplementary Figure 1:**

(A) mRNA relative expression of HMGB1 and PtdSer in NXS2 and Neuro2a NBL cell lines. (B) Supernatant was collected and analyzed for IFNγ and TNF-α release by ELISA with samples in triplicate and experiments repeated 2 times. IFNγ and TNF-α concentration are shown for IL-15 NK cells, 15-4P stimulated NK cells with isotype antibody or anti-Tim-3 blockade. Concentration was calculated based on 5 parameter curve fit interpolated from standard curve. Individual values are plotted with mean and SEM. Two-sided two-sample t tests were performed for comparison between 2 groups. Comparisons between 3 or more groups was analyzed with Brown-Forsythe and Welch one-way ANOVA test with Dunnett's T3 multiple comparisons test (* $p < 0.05$, ** $p < 0.01$, ns = not significant).

**Supplementary Figure 2:**

15-4P stimulated NK cells with isotype control (blue) or anti-TIM-3 blockade (red) were plated with NXS2 NBL at an E:T ratio of 10:1 for 4 hours then NK cells were collected and analyzed by FACs. Percentage (left) and MFI (right) on NK cells are shown for (A) NKG2D and NKp46 percentage and expression on NK cells are shown for all groups. (B) TRAIL, Fas-L, granzyme B, and perforin percent (top) and MFI (bottom) are shown for 15-4P stimulated NK cells and 15-4P stimulated NK cells with anti-TIM-3 antibody. (C) CD107a percentage and expression on 15-4P stimulated NK cells is shown. (D) EOMES, and T-bet percentage and expression, and ratio of NK cells expressing EOMES relative to T-bet for 15-4P stimulated NK cells is shown. Mean with SEM shown for technical replicates (n = 5 per group). Comparisons between 2 groups were analyzed with two-sided two-sample t tests (* $p < 0.05$, ns = not significant).

**Supplementary Figure 3:**

(A) Overall survival and (B) tumor volume for mice receiving allo-HCT and no treatment or treatment with TIM-3 blockade are shown (n = 5 mice per group). Tumor volume was calculated by as V = ½ (Length × Width2). Bars and points with error bars show mean with SEM.

**Supplementary Table 1.**

Expression of differentially expressed genes in 15-4P stimulated NK cells treated with anti-TIM-3 in comparison with 15-4P stimulated NK cells treated with isotype control.

**Supplemental Figure 1.**

**A.**

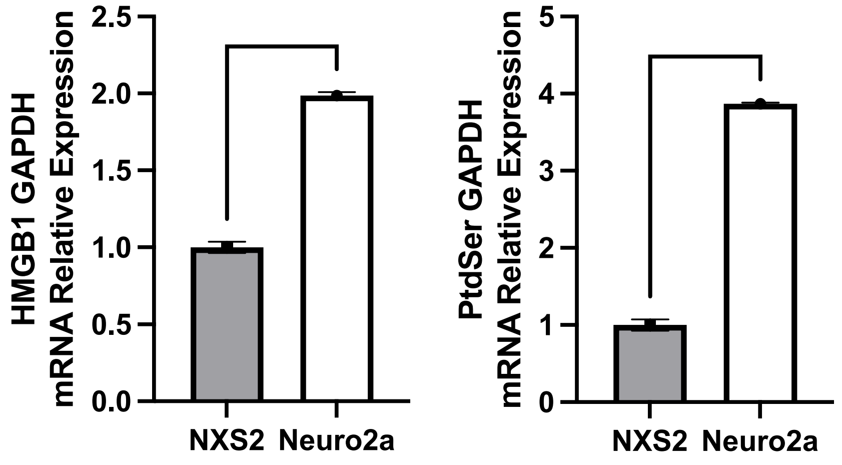


**B.**

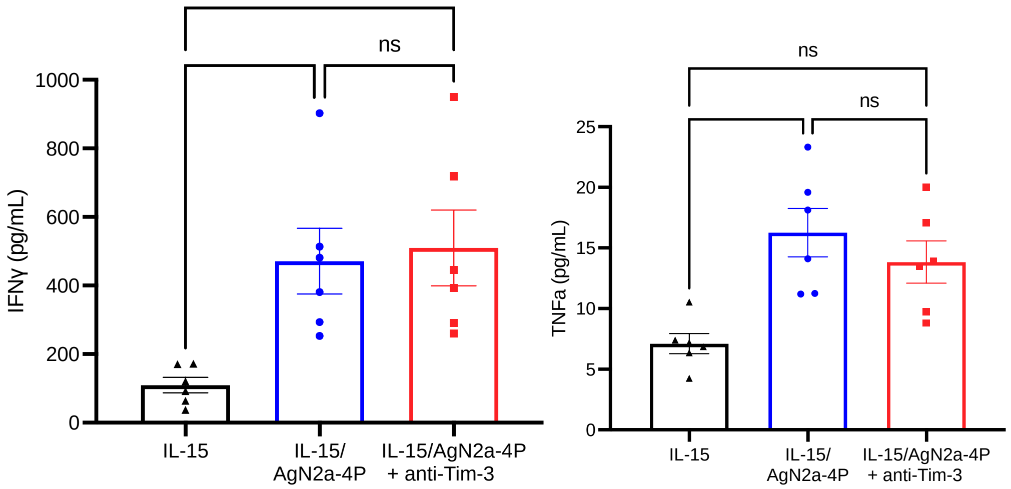

**Supplemental Figure 2.**

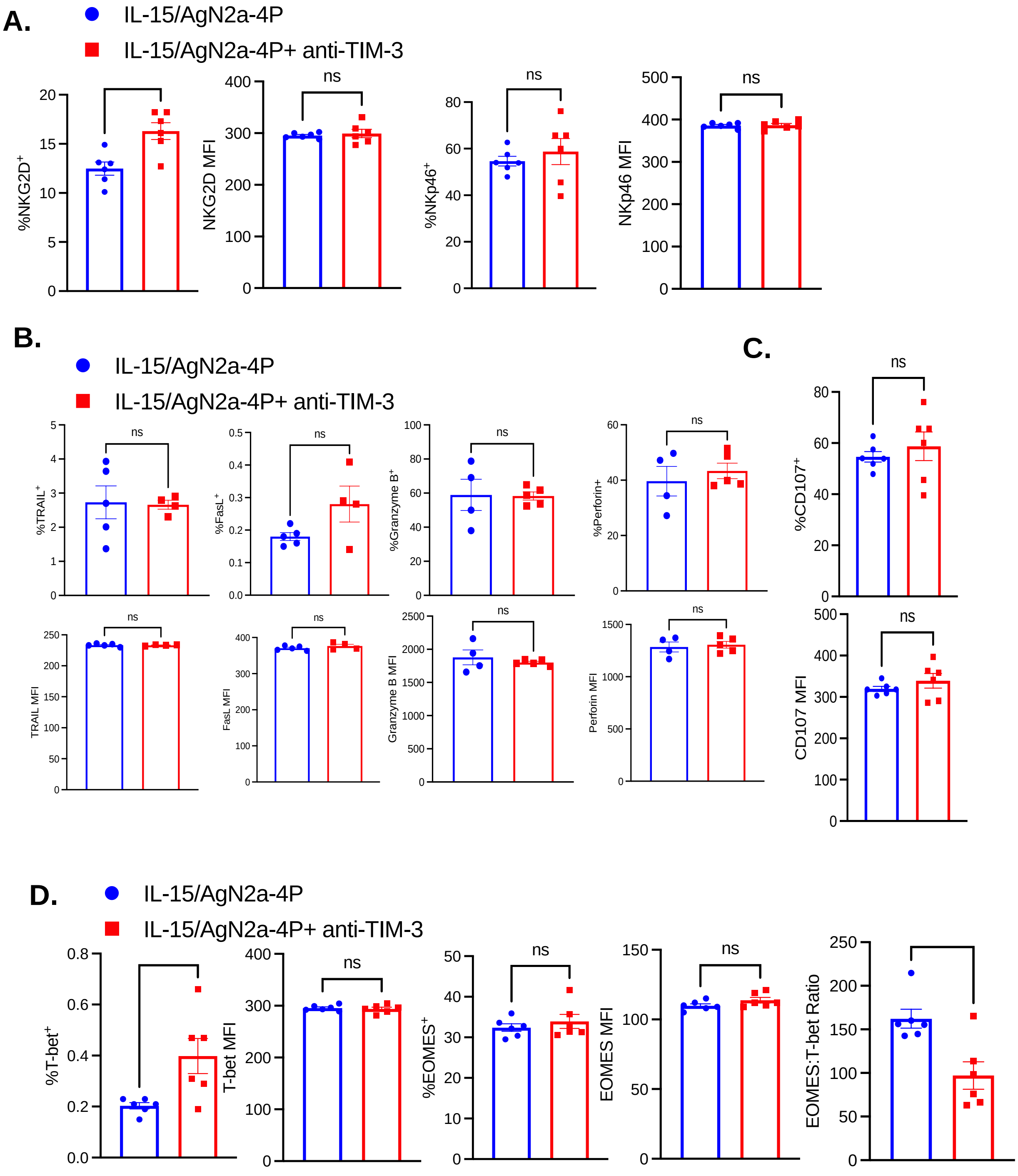

**Supplemental Figure 3**

**A.**

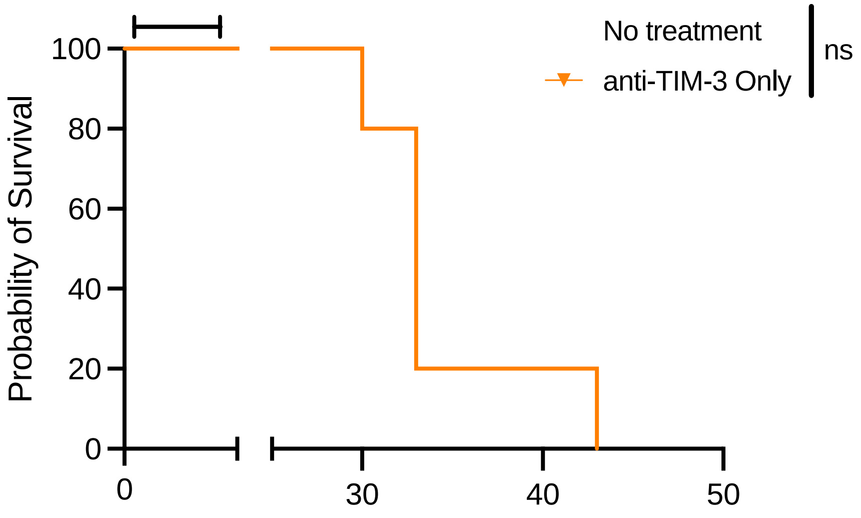


**B.**

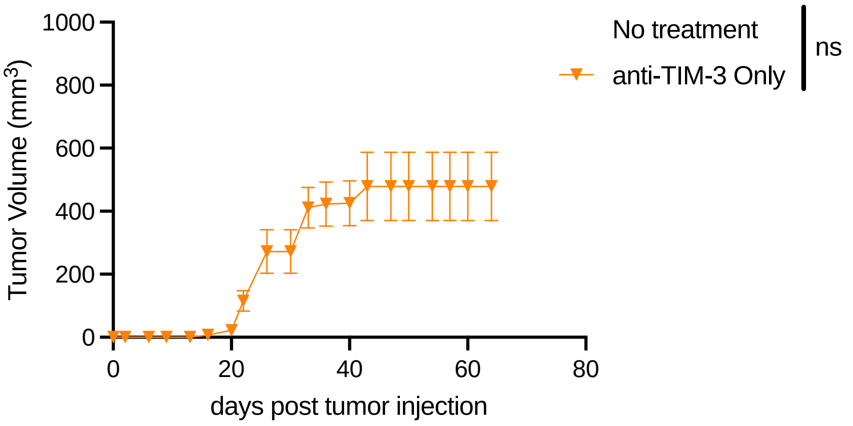


.

#### Supplemental Table 1.

| Gene | Log fold-change | False discovery rate -adjusted p value | Gene name |
|---|---|---|---|
| Icosl | 2.6 | 0.00031 | C-C motif chemokine ligand 1 |
| Tnf | 2.7 | 0.00048 | C-C motif chemokine ligand 3 |
| Ltb | 3.7 | 0.0011 | CD160 antigen |
| Itga2 | 2 | 0.0011 | CD55 molecule, decay accelerating factor for complement |
| Ccl3 | 3.3 | 0.0017 | chymase 1, mast cell |
| Csf2 | 2.8 | 0.0017 | colony stimulating factor 2 (granulocyte-macrophage) |
| Tnfsf8 | 2.7 | 0.0017 | cathepsin G |
| Cd160 | 2.6 | 0.0017 | C-X-C motif chemokine receptor 6 |
| Nfkbia | 2.1 | 0.0017 | fucosyltransferase 7 |
| Socs1 | 2 | 0.002 | guanylate binding protein 5 |
| Gbp5 | 2.5 | 0.0023 | hepatitis A virus cellular receptor 2 |
| Tnfsf14 | 2.6 | 0.0024 | intercellular adhesion molecule 1 |
| Tnfsf10 | 2.4 | 0.0024 | icos ligand |
| Relb | 2 | 0.0024 | interferon induced transmembrane protein 1 |
| Icam1 | 2.4 | 0.003 | interferon induced transmembrane protein 2 |
| Traf3 | 2.1 | 0.003 | interleukin 15 |
| Klrb1 | 2 | 0.0033 | interleukin 21 receptor |
| Cxcr6 | 2.3 | 0.0038 | interferon regulatory factor 8 |
| Il15 | 2 | 0.0038 | immunity-related GTPase family M member 2 |
| Rora | 2.6 | 0.0045 | integrin alpha 2 |
| Xcl1 | 2.6 | 0.0056 | killer cell lectin-like receptor subfamily B member 1 |
| Lta | 2.6 | 0.006 | killer cell lectin-like receptor subfamily G, member 1 |
| Irgm2 | 2.6 | 0.0073 | lymphotoxin A |
| Ccl1 | 3.4 | 0.0077 | lymphotoxin B |
| Il21r | 2.3 | 0.0088 | macrophage migration inhibitory factor (glycosylation-inhibiting factor) |
| Mif | -2.6 | 0.00031 | nuclear factor of kappa light polypeptide gene enhancer in B cells inhibitor, alpha |
| Vegfa | -4 | 0.0012 | perforin 1 (pore forming protein) |
| Havcr2 | -4.4 | 0.0017 | avian reticuloendotheliosis viral (v-rel) oncogene related B |
| Ifitm2 | -4.4 | 0.0024 | RAR-related orphan receptor alpha |
| Ctsg | -5.6 | 0.0024 | suppressor of cytokine signaling 1 |
| Klrg1 | -3.4 | 0.0028 | tumor necrosis factor |
| Irf8 | -2.6 | 0.003 | tumor necrosis factor (ligand) superfamily, member 10 |
| Prf1 | -2.4 | 0.0044 | tumor necrosis factor (ligand) superfamily, member 14 |
| Cd55 | -5 | 0.0044 | tumor necrosis factor (ligand) superfamily, member 8 |
| Ifitm1 | -5.7 | 0.0044 | TNF receptor-associated factor 3 |
| Cma1 | -5 | 0.0056 | vascular endothelial growth factor A |
| Fut7 | -2.3 | 0.006 | chemokine (C motif) ligand 1 |
