## Supplemental Methods for "TIM-3 blockade enhances ex vivo stimulated allogeneic NK cell therapy for relapsed murine neuroblastoma after hematopoietic cell transplant"

### Maintenance of tumor cell lines

Murine NBL Neuro2a cells (H-2a) were obtained from the American Type Culture Collection (ATCC, Manassas, VA). NXS2 is a murine GD2$^{+}$ NBL cell line on an H-2$^{a}$ background and was obtained from Ralph Reisfeld (Scripps Research Institute). Murine NBL AgN2a is an aggressive clone of Neuro2A from which the AgN2a-4P cell line was generated [1]. AgN2a-4P transfectants express CD54, CD80, CD86, and CD137L, and was received as a gift from Dr. Bryon Johnson at the Medical College of Wisconsin (Milwaukee, WI). All tumor cells were maintained in RPMI (Sigma Aldrich, St. Louis, MO) containing 10% fetal bovine serum (FBS) (Sigma Aldrich), Minimal Essential Medium (Thermo Fisher Scientific, Waltham MA), Sodium Pyruvate (Life Technologies, Inc., Gaithersburg, MD), 100 units/mL penicillin (Life Technologies, Inc.) 100units/mL streptomycin (Life Technologies, Inc.), and 1% L-glutamine (Thermo Fisher Scientific). Cell authentication was performed using short tandem repeat analysis (Idexx BioAnalytics, Columbia, MO) and per ATCC guidelines using morphology, growth curves, and Mycoplasma testing within 6 months of use with the MycoStrip Mycoplasma Detection Kit (Invitrogen, Waltham, MA). Cells were maintained at 37°C in 5% $CO_2$ and used after 3-5 passages in culture after thawing.

### Incucyte characterization of NK cell cytotoxicity

Murine NBL cell lines Neuro-2A or NXS2 were plated with day 12 expanded IL-15 NK cells or IL-15/AgN2a-4P NK cells at an effector to target (E: T) ratio 10:1 (100,000 Effector cells: 10,000 target cells) (n = 3 - 6 wells per condition). For antibody blockade, cells were incubated with 10 ug/mL purified rat anti-mouse anti-TIM-3 antibody (Bio X Cell Cat. No. BE0115, Lebanon, NH) or purified rat anti-mouse anti-IgG2a (Bio X Cell Cat. No. BE0089) antibody as a negative control for 30 minutes at 37°C prior to use. Target cells were incubated with 1 uM Staurosporine for the maximum killing positive control. To detect apoptotic cell death, 5 uM NucView488 Caspase-3 (Biotium Fremont, CA cat. No. 4440) was added to each well in accordance with the manufacturer's protocol. To track mitochondrial membrane potential in tumor targets, Neuro2a or NXS2 cells were prelabeled with MitoView™ 633 dye (Biotium Fremont, CA cat. No. 70055-T). Effectors and targets were cocultured in the IncuCyte S3 System (Sartorius AG, Göttingen, Germany) live cell imaging platform at 10X magnification with images taken every 4 hours for 24 hours. NK cell-driven target cell lysis was analyzed using IncuCyte analysis software (Sartorius) by size exclusion and fluorescence total object area (μm2/well) to evaluate the number of target cells present undergoing apoptosis.

**Phenotypic characterization and analysis with flow cytometry**

To analyze the tumor microenvironment, murine tumor cell lines Neuro2a and NXS2 were prepared at a concentration of 1 x $10^6$ in a single cell suspension and stained with CEACAM-1 PE-Cy7 (Biolegend Cat. No. 134516), Galectin-9 APC (Biolegend Cat. No. 136110 ), MHC I (H-2) FITC (Biolegend Cat. No. 125508), ULBP-1/MULT-1 PE (R&D Systems Cat. No. FAB2588P). Cells were then washed in FACS buffer containing PBS (Corning) with 0.2% FBS (Sigma Aldrich) and 0.1% sodium azide (Sigma Aldrich). Live/dead Staining was performed using Ghost Dye Red 780 (13-0865-T500, Tonbo Biosciences, San Diego, CA).

In NK cell characterization experiments, murine B6 NK cells were prepared at a concentration of 1x$10^6$ in a single cell suspension. Briefly, cells were washed with PBS, incubated with mouse TruStain FcX Receptor Blocking Antibody (Biolegend Cat. No. Cat# 101320, San Diego, CA), then stained with the following fluorophore-conjugated monoclonal antibodies at 4°C for 30 minutes; CD3 Pacific Blue (Biolegend Cat. No. 100214), NK1.1 BV510 (Biolegend Cat. No. 108738), NKG2D PE-Dazzle 594 (Biolegend Cat. No. 130213), NKp46 FITC (Biolegend Cat. No. 137606), TIM-3 PE (Biolegend Cat. No. 119704), and GhostRed780 viability dye (Tonbo Biosciences Cat. No. 13-0865-T100) in panel 1, and CD3 Pacific Blue (Biolegend Cat. No. 100214), NK1.1 BV510 (Biolegend Cat. No. 108738), Fas-L PE (Biolegend Cat. No. 106606), and CD253 (TRAIL) PerCP-Cy5.5 (Biolegend Cat. No. 142108) and GhostRed780 viability dye (Tonbo Biosciences Cat. No. 13-0865-T100) in panel 2.

For proliferation experiments, cells were stained with GhostRed780 viability dye (Tonbo Biosciences Cat. No. 13-0865-T100), fixed with 70% ethanol and incubated at -20°C overnight. Cells were then stained with Ki-67 PerCP-eFluor 710 (Invitrogen Cat. No. 46-5698-82).

In all flow cytometry experiments, cells were analyzed using the Attune NxT Cytometer (Invitrogen) and FCS files were analyzed using FlowJo software version 10 (Ashland, OR). Doublets were excluded by forward scatter and side scatter singlets and gated on live cells. B6 NK cells were defined by $CD3^-$, $NK1.1^+$ and median fluorescence intensity was measured to assess expression level for activation markers.

**Degranulation and cytokine production**

Murine B6 IL-15 NK cells and B6 IL-15/AgN2a-4P NK cells were plated with Neuro2a or NXS2 murine NBL in 96 well flat bottom plates at effector to target (E: T) ratio of 10:1 with target cell seeding density at 10,000 cells/well and effect cell density at 100,000 cells/well (n = 5 wells per condition). For antibody blockade, cells were incubated with 10 ug/mL purified rat anti-mouse anti-TIM-3 (Bio X Cell Cat. No. BE0115, Lebanon, NH) or purified rat anti-mouse anti-IgG2a (Bio X Cell Cat. No. BE0089) as a negative control for 30 minutes at 37°C prior to use. After incubation for 30 minutes at 37°C, GolgiSTOP (monesin) (BD Biosciences Cat. No. 554724, Franklin Lakes, NJ) and GolgiPLUG (brefeldin A) (BD Biosciences Cat. No. 555029) were added to each well, and the cells were incubated for another 4 hours at 37°C. For degranulation and other flow cytometry analysis, cells were harvested and stained. For intracellular cytokines, cells were fixed using BD CytoFix buffer and permeabilized with BD Perm/Wash buffer. Cells were stained with Granzyme B PE-Cy7 (Biolegend Cat. No. 372214), Perforin APC (Biolegend Cat. No. 154304), T-bet BV605 (Biolegend Cat. No. 644817), EOMES AF700 (Biolegend Cat. No. 157703), CD107a BV711 (Biolegend Cat. No. 121631) and GhostRed780 viability dye (Tonbo Biosciences Cat. No. 13-0865-T100). Cells were analyzed using the Attune NxT Cytometer (Invitrogen) and FCS files were analyzed using FlowJo software version 10 (Ashland, OR). Doublets were excluded by forward scatter and side scatter singlets and gated on live cells. B6 NK cells were defined by $CD3^-$, $NK1.1^+$ and median fluorescence intensity was measured to assess expression level for activation markers.

To assess TNFα and IFNγ production, supernatant was collected and frozen at -80°C for later analysis by ELISA. Collected undiluted cell supernatant was assayed for IFNγ detection using Mouse IFNγ ELISA MAX (Biolegend Cat. No. 430815) or TNFα detection using Mouse TNFα (Biolegend Cat. No. 430904). ELISA plates were read using a CLARIOstar microplate reader (BMG Labtech, Cary, NC) at 450 nm wavelength. Analyte concentration was determined based on a non-linear 5-parameter logistic curve calculated using Prism 9 (Graphpad Software, San Diego, CA).

**RNA isolation, reverse transcription, and RT-qPCR.**

Prior to RNA isolation, IL-15/AgN2a-4P expanded B6 NK cells were stained and purified using an NK cell isolation kit (Miltenyi Biotec Cat. No 130-115-818) on the AutoMACs Pro (Miltenyi Biotec) according to manufacturer's protocol. Total RNA was isolated from NK cells or primary tumor tissue using the RNeasy Mini Kit (QIAGEN) following the manufacturer's protocol. RNA concentrations were quantified on NanoDrop 1000 Spectrophotometer (Thermo Fisher Scientific), and absorbance was measured at 260/280 (high purity defined as ratio ~2.0) and 260/230 (high purity defined as ratio 2.0-2.2) to assess the purity of isolated mRNA. cDNA was synthesized using iScript Reverse Transcription Supermix (Bio-Rad Laboratories, Inc., Hercules, CA). mRNA expression was detected using qRT-PCR using specific primers for human, and mouse expression (supplemental table) on the Veriti 96-Well PCR Thermal Cycler (Applied Biosystems Waltham, MA).

**RNA sequencing and gene expression analysis**

NK cells were harvested at day 12 and exposed to anti-TIM-3 or IgG2a isotype control antibody for 4 hours. Prior to RNA isolation, NK cells (n = 2) were purified using the AutoMACS pro-separator (Miltenyi Biotec, San Jose, CA). Total RNA was isolated from purified NK cells using an RNA isolation kit (Qiagen, Germantown, MD, USA) according to the manufacturer's protocol. RNA was subjected to quality control assessments for A260/230, A260/280, and DV200 and Gene expression was analyzed with the using the nCounter Mouse PanCancer Immune Profiling Panel Kit (XT_PGX_MmV1_CancerImm_CSO) (NanoString Technologies, Inc., Seattle, WA, USA) (Supplementary Table S3). The NanoTube Bioconductor package was used for processing, quality control according to standard nSolver steps, normalization, and analysis of raw nCounter expression data. Differential expression gene (DEG) analysis and group-vs-group comparisons were generated using the limma R package [2]. Differentially expressed genes were identified using thresholds of log fold-change (FC) of $\geq 2$ and a q value of $\leq 0.05$.

**In vivo depletion procedure**

For NK cell depletion experiments, BMCs were processed, T cell depleted as previously described, and depleted of NK cells using biotin conjugated NK1.1 antibody (Biolegend Cat. No. 108703), and anti-biotin microbeads (Miltenyi Biotec Cat. No. 130-090-485). Irradiated B6AJF1 recipient mice received retroorbital injection of 5 x 106 donor-derived NK depleted BMCs and 1x106 T cells. Additionally, mice received of 200μg of anti-NK1.1 (Bio X Cell Cat. No. BP0036) by intraperitoneal injection every 4 days to maintain NK depletion. For T cell depletion experiments BMCs were processed, T cell depleted as previously described, and irradiated B6AJF1 recipient mice received retroorbital injection of 5x106 donor-derived BM cells without T cell add-back. Survival after transplant was monitored biweekly. Mice were randomized, assigned to groups, and assessed for presentation of clinical GVHD by using a scoring system which sums the following five parameters: percent weight loss, posture, activity, fur texture and skin integrity [3].
